# *In vitro* and *in vivo* NIR Fluorescence Lifetime Imaging with a time-gated SPAD camera

**DOI:** 10.1101/2021.12.26.474189

**Authors:** Jason T. Smith, Alena Rudkouskaya, Shan Gao, Juhi M. Gupta, Arin Ulku, Claudio Bruschini, Edoardo Charbon, Shimon Weiss, Margarida Barroso, Xavier Intes, Xavier Michalet

## Abstract

Near-infrared (NIR) fluorescence lifetime imaging (FLI) provides a unique contrast mechanism to monitor biological parameters and molecular events *in vivo*. Single-photon avalanche photodiode (SPAD) cameras have been recently demonstrated in FLI microscopy (FLIM) applications, but their suitability for *in vivo* macroscopic FLI (MFLI) in deep tissues remains to be demonstrated. Herein, we report *in vivo* NIR MFLI measurement with SwissSPAD2, a large time-gated SPAD camera. We first benchmark its performance in well-controlled *in vitro* experiments, ranging from monitoring environmental effects on fluorescence lifetime, to quantifying Förster Resonant Energy Transfer (FRET) between dyes. Next, we use it for *in vivo* studies of target-drug engagement in live and intact tumor xenografts using FRET. Information obtained with SwissSPAD2 was successfully compared to that obtained with a gated-ICCD camera, using two different approaches. Our results demonstrate that SPAD cameras offer a powerful technology for *in vivo* preclinical applications in the NIR window.

## I. Introduction

Preclinical molecular imaging is used in early drug development^1,2^ and as a research tool to better understand the biology of drug resistance. Two imaging techniques provide the high sensitivity needed to detect biomarkers during and after drug delivery: nuclear (PET) and optical imaging. They both allow non-invasive assessment of delivery efficacy, pharmacokinetics, and response in longitudinal studies^1–3^. PET is good at quantitative molecular imaging of targeted drug delivery in live subjects^4^, providing spatial and temporal distribution of labeled probes in living animals^4^, but is limited to a single targeted radiotracer. Co-localization of a radiotracer-labeled antibody-drug conjugates with the pathological site unfortunately does not provide unequivocal evidence of actual binding to the target protein i.e. receptor engagement, which is essential to elicit the cellular response necessary to kill cancer cells^56–8^. By contrast, optical imaging methods^9,10^ and in particular fluorescence imaging^11,12^, which offer the possibility to monitor several probes simultaneously, can lift this ambiguity. Fluorescence lifetime imaging (FLI) can additionally report on numerous intracellular parameters such as metabolic status^13^, reactive oxygen species^14^ and intracellular pH^15^ or to quantify Förster Resonant Energy Transfer (FRET), a powerful technique used to study protein-protein interactions and biosensor activity^16^.

We use macroscopic FLI-FRET (MFLI-FRET) to quantify *in vivo* drug-target engagement in live, intact animals over large fields of view^17–19^: the reduction of donor fluorophore probe lifetime upon drug-target engagement results from the proximity of an another acceptor fluorophore-labeled probe bound to the same target. Donor lifetime can be measured with high sensitivity and dispense with corrections required in other FRET techniques, such as sensitized-emission FRET^20^. On the other hand, MFLI-FRET data acquisition requires more complex instrumentation compared to intensity-based fluorescence imaging. Specifically, time-gated ICCD cameras, which are the detectors of choice for MFLI applications, are expensive, prone to photocathode degradation, susceptible to damage from overexposure, and as a dated technology, have limited room for technical improvements. By contrast, time-resolved CMOS SPAD arrays (SPAD cameras) have undergone tremendous developments over the past decade, and are poised to become a competitive solution for fluorescence lifetime imaging as discussed here^21^.

SwissSPAD2 (SS2) is a very large time-resolved SPAD camera with single-photon sensitivity, developed specifically for FLI applications^22–24^. This time-gated imaging sensor is comprised of 512 512 SPAD pixels, each associated with a 1-bit memory, readout as a whole at up to 97,700 frames per second^22^. SS2’s capabilities for FLIM applications in the visible spectrum have been recently described^22–24^. Although these studies demonstrated SS2’s potential for microscopic biological applications, there are outstanding challenges involved with using it for pre-clinical macroscopic imaging applications. First, NIR dyes used for *in vivo* small animal imaging are challenging to detect, due to the reduced photon detection probability of silicon SPADs in the NIR. Next, most NIR dyes exhibit lifetimes far shorter than visible dyes (few hundreds of picoseconds – ps – compared to few nanoseconds). Lastly, MFLI-FRET involves quantifying fluorescence decays from two or more fluorophore species or states simultaneously, which requires larger signal-to-noise compared to mono-exponential cases, leading to additional challenges^25^.

Herein, we report the first application of SS2 in a variety of MFLI measurements of NIR fluorescent samples *in vitro* and *in vivo*, systematically comparing it to a state-of-the-art gated-ICCD (Supplementary Fig. S1). First, we study the sub-nanosecond lifetime NIR dye Alexa Fluor 750 with both detectors, using two distinct methods: nonlinear least square fit (NLSF) and phasor analysis. We then quantify the lifetime of the clinically relevant NIR dye IRDye 800CW^26^ as a function of molecular microenvironment. MFLI measurements of NIR-FRET pairs in different ratios characterized by multi-exponential decays conclude these *in vitro* benchmarks. Next, we use SS2 for noninvasive preclinical MFLI-FRET imaging of live mice carrying HER2 positive tumor models. Two clinical cancer drugs, the anti-HER2 monoclonal antibody (mAb) Trastuzumab (TZM) and the anti-EGFR mAb Cetuximab (CTM), both labeled with NIR donor and acceptor dyes are used as FRET probes to visualize HER2-and EGFR-positive tumors in live mice. We successfully characterize organs of interest across the mouse body based on lifetime information, and support these characteristics by a systematic comparison between detectors (SS2 and ICCD) and methods (NLSF and phasor analysis), demonstrating the suitability of SS2 for these challenging *in vitro* and *in vivo* applications.

## II. RESULTS

### NIR-MFLI lifetime measurement as a function of molecular environment (IRDye 800CW-2DG)

To assess SS2’s ability to resolve minute differences in NIR lifetimes, such as those expected for fluorescent dyes exposed to different microenvironments, solutions of IRDye 800CW conjugated to 2-deoxyglucose (2DG) were prepared in aqueous buffers with different pH, as well as in DMSO. IRDye 800CW-2DG is typically used as a reporter of metabolic activity in small animal model. Although no systematic study of environmental effects on IRDye 800CW has been published, there is evidence of pH effects on fluorescence quantum yield, and hence radiative lifetime, in other cyanine dyes; for instance, some ICG derivatives exhibit higher quantum yields and lifetimes at lower pH^27^. Similarly, DMSO has been observed to increase the quantum yield/radiative lifetime of ICG, among other indocyanine derivatives^28^. We thus performed MFLI studies of solutions of IRDye 800CW-2DG in various environments with both SS2 and ICCD, and compared the resulting lifetimes obtained with either NLSF or phasor analysis.

Nonlinear Least Square Fit (NLSF, see *Online Methods*) is a standard technique which only requires an instrument response function (IRF) measurement, as well as good signal to noise ratio (SNR) to obtain accurate results, but can be computationally demanding^25,29^. A user-friendly and fit-free alternative to NLSF analysis is the phasor approach^18,30–3434^, which is based on the computation of a pair of Fourier coefficients of the decay (Fourier harmonic *n* or phasor frequency *f = n/T*), which are then represented as a point in the *phasor plot*. In the case of species characterized by mono-exponential decays, phasors are located on the *universal circle* (UC) and their lifetime can be retrieved geometrically (referred to as the *phase lifetime*, see *Online Methods*). Phasor analysis of *time-gated* decays proceeds similarly, with the universal circle replaced by a modified curve^36^ (dubbed *single-exponential phasor locus* or SEPL following ref. ^34^), which is barely distinguishable from the UC as the resolution (number of gates) increases. Other differences can appear in the case of (i) incomplete or truncated decays (as is the case for the ICCD data obtained in this study), where a different phasor frequency *f =* 1*/D* (decay support window length *D* < *T*) is preferable, or (ii) gate with non-ideal shapes (which characterize both detectors used here)^34^. Details on the best way to handle these differences can be found in ref. ^34^ and are summarized in the *Online Methods*.

We first verified that both methods retrieve lifetimes accurately independently from signal intensity by analyzing solutions of a single NIR dye prepared at different concentration (see Supplementary Note 1 and Fig. S2).

### NLSF Analysis

As shown for the AF750 measurements (Fig. S2), intensity maps obtained with both ICCD and SS2 for IRDye 800CW (Fig. 1A & B, respectively) are comparable despite setup differences. However, very distinct normalized decays are observable in some wells, indicative of different lifetimes (Fig. 1C and 1F for the ICCD and SS2, respectively). Lifetime results obtained for both imagers (Fig. 1E,H) indicate no variation of IRDye 800CW’s lifetime (*τ* = 0.5 ns) as function of pH in the range 5.5 to 7.5, but a measurable increase at lower pH (*τ* = 0.6 ns at pH 4.5), which is also the pH of our distilled water (H_2_O sample). Interestingly, the lifetime in DMSO more than doubles (*τ* = 1.1 ns) compared to the values in aqueous buffers, likely reflecting the increased orientation polarizability of DMSO, as observed for similar dyes^37,38^. As observed before, both detectors provide essentially identical results (Fig. 1O), with a maximum relative difference between the two measurements of 2.7%.

**Fig. 1:**
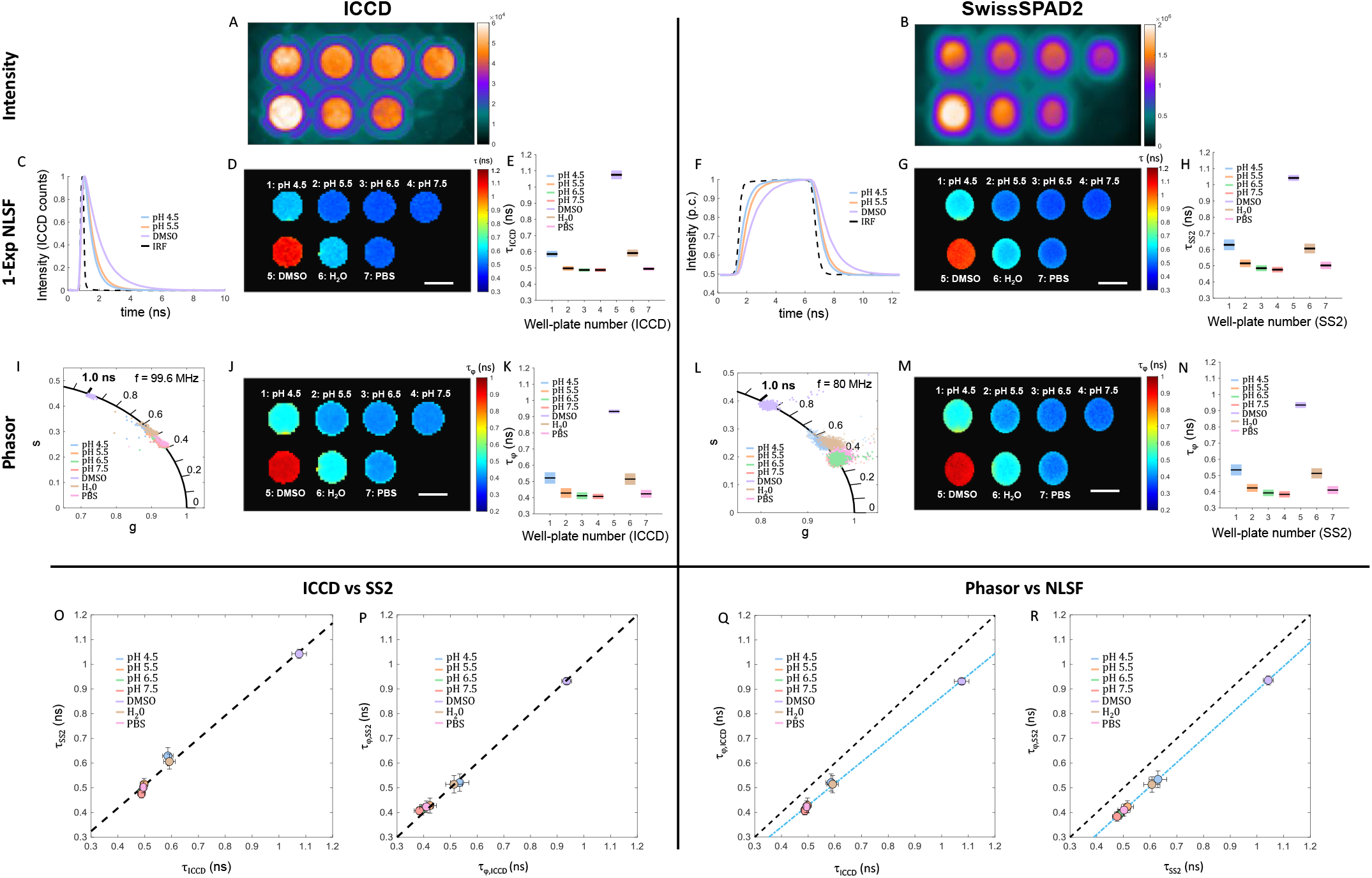
Environment effects on IRDye 800CW-2DG. A, B: Fluorescence intensity images. A, ICCD: MCP voltage = 400 V, integration time = 308 ms, illumination power = 0.76 mW/cm^2^. B, SS2: integration time = 1.02 s, illumination power = 1.53 mW/cm^2^; All wells were prepared at constant fluorophore concentration (15 μM) and are labeled with their respective buffer/pH. C, F: Representative ICCD and SS2 normalized single-pixel decays for 3 of the wells, plotted with the corresponding IRF. D, G: lifetime maps obtained by NLSF. E, H: Boxplot summarizing lifetime results for all wells. J, M: Phasor plot for each well and K, N: pixel-wise phase lifetime maps obtained with the ICCD and SS2 detectors. L, O: Boxplot quantifying phase lifetime results for all wells. O: Scatter plot of averaged lifetime results for ICCD and SS2. P: Scatter plot of SS2 phase lifetime versus ICCD phase lifetime. Q, R: Scatter plot of averaged NLSF lifetime results versus averaged phase lifetime results for both ICCD and SS2, respectively. Error bars represent 1-standard deviation in O-R. Scale bar in A, B, D, G, J, M: 3 mm.

### Phasor analysis

Phasor plots for all IRDye 800CW-2DG mixtures (Fig. 1I,L), as well as corresponding pixel-wise phase lifetime maps (Fig. 1J,M), are shown in Fig. 1 for the ICCD and SS2. The phase lifetime results obtained with both cameras are in excellent qualitative and quantitative agreement with one another (Fig. 1P). Comparison between phase lifetime (*τ*_φ_) and NLSF lifetime (*τ*) for each well shows a small positive bias (<100 ps) for NLSF lifetimes (Fig. 1Q,R). Because it is common to both cameras, this suggests that the IRF data used for convolution (NLSF) and calibration (phasor analysis) might not be perfectly adequate (see *Online Methods*). In any case, once characterized, this systematic bias can easily be corrected for post-analysis and will be neglected in the remainder of this study.

### *In vitro* NIR MFLI-FRET measurements: IgG-AF700/anti-IgG-AF750 mixtures

FRET between neighboring fluorescent dyes having overlapping emission (donor molecule) and absorption spectra (acceptor molecule) is widely used as a molecular ruler^39^ but also as a qualitative reporter of protein-protein interactions. FRET acts as a proximity assay thanks to its nanoscale range, set by the Förster radius *R*_*0*_, of the order of a few nm, which depends on the donor and acceptor photophysical properties. One of the characteristic signatures of FRET is a decrease of the donor fluorescence lifetime, *τ*_*D*_, in the presence of a nearby acceptor molecule (*τ*_*DA*_ < *τ*_*D*_). In an ideal mixture of interacting donor- and acceptor-labeled molecules, donor molecules are either engaged in FRET with an acceptor molecule at a certain distance (FRET fraction: *f*_*FRET*_, lifetime: *τ*_*DA*_) or isolated (donor-only fraction: 1-*f*_*FRET*_, lifetime: *τ*_*D*_), resulting in an observed bi-exponential fluorescence decay. The FRET fraction *f*_*FRET*_ can then be quantified by a fit of the measured decay to a bi-exponential model, or indirectly, by measuring the average lifetime. Increasing FRET fraction decreases the effective donor lifetime, measured as either the average donor lifetime or approximated as the best single-exponential decay fit^40,41^.

AF700/AF750 is a NIR FRET pair suitable for *in vivo* optical imaging applications, which we have used to assess transferrin^17–19^ and TZM^42,43^ tumor delivery and efficacy in small animal models. To verify SS2’s ability to quantify FRET, a controlled *in vitro* NIR MFLI-FRET experiment was first carried out. A multiwell-plate was prepared using the AF700/AF750 FRET pair (Förster radius *R*_*0*_ ∼8 nm^44^), with each dye conjugated to a complementary IgG and anti-IgG pair (IgG-AF700 and anti-IgG-AF750 respectively). Each well is characterized by a different acceptor-to-donor ratio (A:D, of the form *n*:1 with *n* = 0 - 3). As before, we acquired data with both detectors using similar conditions, and processed data using both NLSF and phasor analysis.

### NLSF analysis

*Single-exponential* NLSF analysis reveals the expected trend of a decreasing donor lifetime with increasing A:D ratio (Fig. S4), as well as close agreement between both detectors. While this agreement is important, single-exponential fits are not sufficient for the quantitative study of FRET mixtures, as there is no simple relationship between fitted single-exponential lifetime *τ* and FRET fraction *f*_*FRET*_. Hence, *bi-exponential* NLSF analysis of the data provides a more accurate interpretation of the data despite relying on some simplifying assumptions. Here, the lifetimes of donor-only (*τ*_1_ = *τ*_D_ = 1 ns) and quenched-donor (*τ*_2_ = 0.265 ns) were retrieved through ROI-level decay fitting (Fig. S5).

Following conventional practices, these values of *τ*_1_ and *τ*_2_ were fixed for all subsequent pixel-level analyses, leaving only 3 free parameters for the NLSF analysis: baseline *B* and amplitudes *A*_*1*_ and *A*_*2*_ (*Online Methods*, Eq. (1)). The FRET intensity fraction *f*_*FRET*_ (*Online Methods*, Eq. (4)) values yield similar results for both imagers (Fig. 2D,G,E,H). Importantly, both results reveal the expected linear increase of the calculated FRET fraction as a function of acceptor-to-donor ratio (Fig. 2O).

**Fig. 2:**
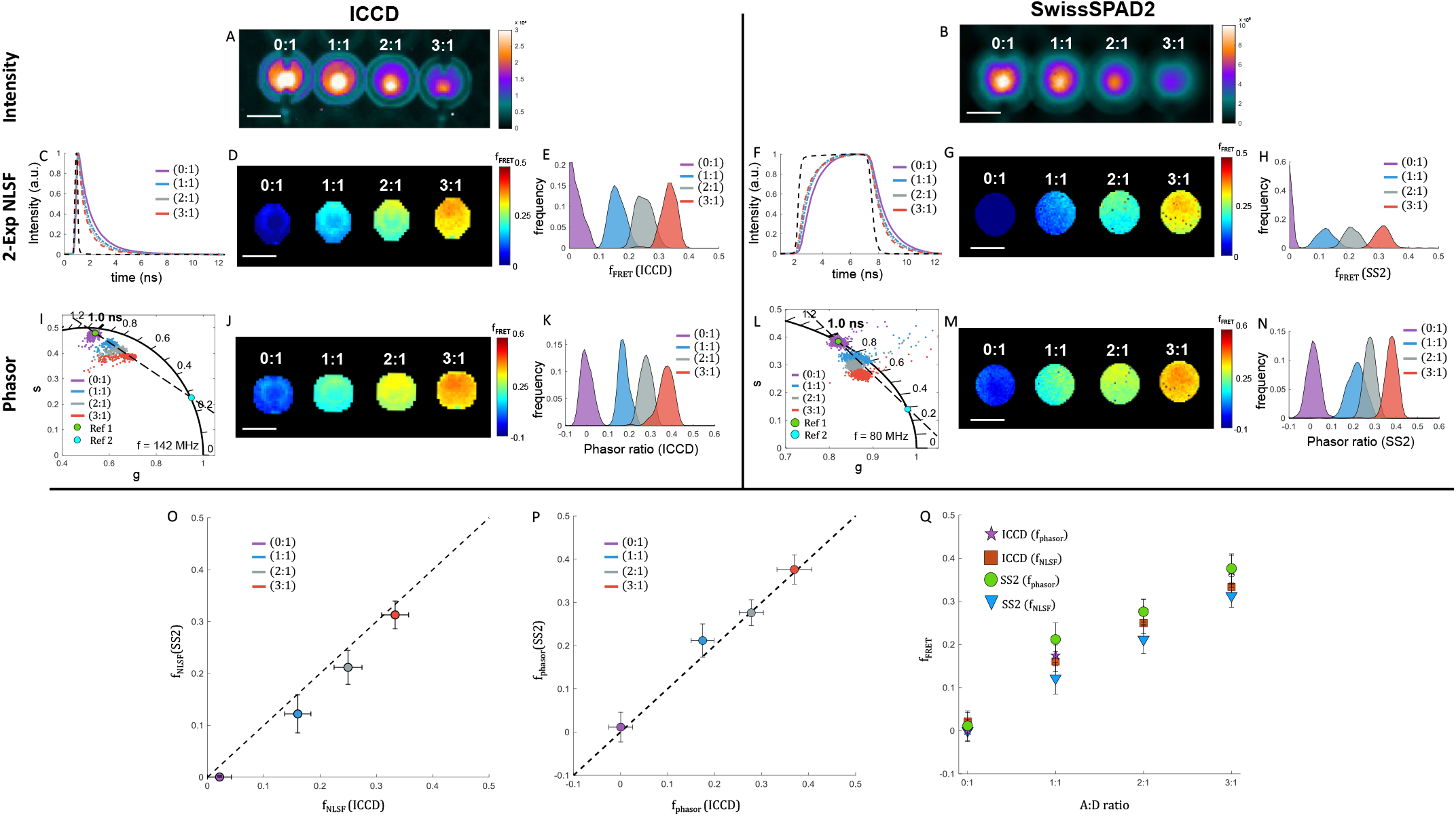
AF700/AF750 FRET pair series. A, B: Fluorescence intensity images. A: ICCD; MCP voltage = 450 V, integration time = 359 ms, illumination power = 1.52 mW/cm^2^. B: SS2; integration time = 4.08 s, illumination power = 2.29 mW/cm^2^; Wells contained solutions of labeled antibodies in PBS buffer with acceptor to donor A:D ratio from 0:1 to 3:1 indicated above each well, with a constant donor fluorophore concentration of 32 μM. C, F: Representative ICCD and SS2 normalized single-pixel decays for the different wells, plotted with the corresponding IRF. D, G: MFLI FRET-fraction maps obtained by bi-exponential NLSF for both cameras. E, H: Corresponding *f*_*FRET*_ KDE distributions for each well obtained for both cameras. I, L: phasor scatter plots overlaid with linear fit (dashed black line) and reference lifetimes set as the centroid of the donor-only well’s cluster (green dot, reference 1) and as the linear fit intersection with the SEPL (blue dot, reference 2) – resulting in *τ*_*1,ICCD*_ = 0.98 ns, *τ*_*2,ICCD*_ = 0.27 ns and *τ*_*1,SS2*_ = 1.01 ns, *τ*_*2,SS2*_ = 0.17 ns. J, M: pixel-wise phasor ratio maps and K, N: phasor-ratio KDE distributions for the different wells (acceptor-donor ratios indicated A:D in parenthesis, color code matching that of panels J, L). O: Scatter plot (mean ± standard deviation) showing the FRET-fraction measured via bi-exponential NLSF for each well with the ICCD versus that measured with SS2. P: Scatter plot of phasor ratio results (mean ± standard deviation) for ICCD and SS2. R: Scatter plot of FRET fractions obtained by NLSF (panels C-H) and phasor ratio results for ICCD and SS2 (panels I-N) as a function of A:D ratio. Scale bar in A, B, D, G, J, M: 3 mm.

### Phasor analysis

Although NSLF analysis as performed above provides important information, it relies on assumptions that may not always be fulfilled. By contrast, phasor analysis of mixtures does not rely on any specific functional form for the reference decays (donor only species or fully quenched donor decay). However, because the decays of mixtures are linear combinations of donor and quenched donor decays, their phasors are also linear combinations of the donor and quenched donor phasors, and are therefore aligned along a segment connecting donor and quenched donor phasors^18,45^.

As hypothesized in the NLSF analysis above, in the ideal case where the donor-only species (reference 1) and the FRET pair species (reference 2) can be assumed to be single-exponential decays, the phasor of each species is located on the SEPL. The relative distance of a particular sample’s phasor with respect to reference 2 (called phasor ratio^18,46^), is equal to the intensity fraction of reference 1 (Eq. (8)). Hence, in that ideal case, the calculated intensity fraction should be equal to that obtained by bi-exponential NLSF analysis of the recorded decay (*Online Methods*, Eq. (4)).

In panels I-N of Fig. 2, we show the result of phasor analysis of the datasets discussed above. The references used for phasor ratio quantification were retrieved by computing the intersection of the SEPL with a line obtained as best fit through phasors from all four wells (Fig. 2I,L).The resulting phasor ratio analysis (Fig. 2J,K,M,N) shows a remarkable agreement for both cameras (Fig. 2P).

FRET fraction results retrieved through both cameras using NLSF and phasor are in good agreement (Fig. 2Q). Noticeably, in the case of phasor ratio analysis, although both datasets are computed with different phasor frequencies and with automatically defined references, both analyses result in reference phasors which are remarkably similar to those chosen for NSLF analysis (donor-only: [*τ*_*1,ICCD*_ = 0.98 ns, *τ*_*1,SS2*_ = 1.01 ns] and quenched donor: [*τ*_*2,ICCD*_ = 0.27 ns and *τ*_*2,SS2*_ = 0.17 ns], respectively).

### *In vivo* NIR MFLI-FRET measurements: Trastuzumab-HER2 engagement in mouse xenografts

Having established the *in vitro* equivalence of time-gated ICCD data and SS2 data, we extended this comparison to *in vivo* small animal experiments. We used xenograft models of human breast cancer (AU565) and ovarian cancer (SK-OV-3) selected due to their HER2 overexpression and distinct challenges regarding effective drug delivery^47^, to compare drug-receptor engagement efficiency across two tumor models. AU565 and SK-OV-3 cells overexpress HER2 receptors, which are targeted by Trastuzumab (TZM), a monoclonal antibody used as anti-HER2 breast cancer therapy. In our approach, AF700 or AF750 are conjugated to TZM and intravenously injected into mice carrying HER2 positive tumor xenografts. Thus, the amount of recorded FRET between the two probes provides a measure of TZM-receptor binding. Noninvasive MFLI-FRET imaging of nude mice bearing AU565 and SK-OV-3 tumor xenografts was performed upon intravenous injection of NIR-labeled TZM pair with both time-resolved detectors in two different mice (see *Online Methods*). Mice are imaged at 24 hr post-injection (p.i.) using the ICCD (mouse 1) or SS2 (mouse 2), followed by two other imaging sessions of mouse 2 at 48 hr (SS2) and 51 hr (ICCD) p.i., respectively.

### NLSF Analysis

NLSF analysis of the 24 hr p.i. data was limited to the two tumors and urinary bladder regions of interests (ROIs) in which significant AF700 signal was observed (Fig. 3A,B). As for the *in vitro* FRET analysis, pixel-wise single-exponential NLSF analysis of the observed decays (Fig. 3C,F) was first performed to obtain a qualitative understanding of the data (Supplementary Fig. S6C-F). The lifetime of AF700-TZM in the urinary bladders is close to 1 ns and distributed uniformly. These results are consistent with the expected detection of donor-only labeled TZM as measured in different pH conditions in Fig. S7. By contrast, AF700-TZM’s lifetime is noticeably shorter in the AU565 tumors, as previously reported^43^, but not in SK-OV-3, suggesting that FRET and thus TZM-HER2 binding is occurring at a higher level in AU565 tumors than SK-OV-3.

**Fig. 3:**
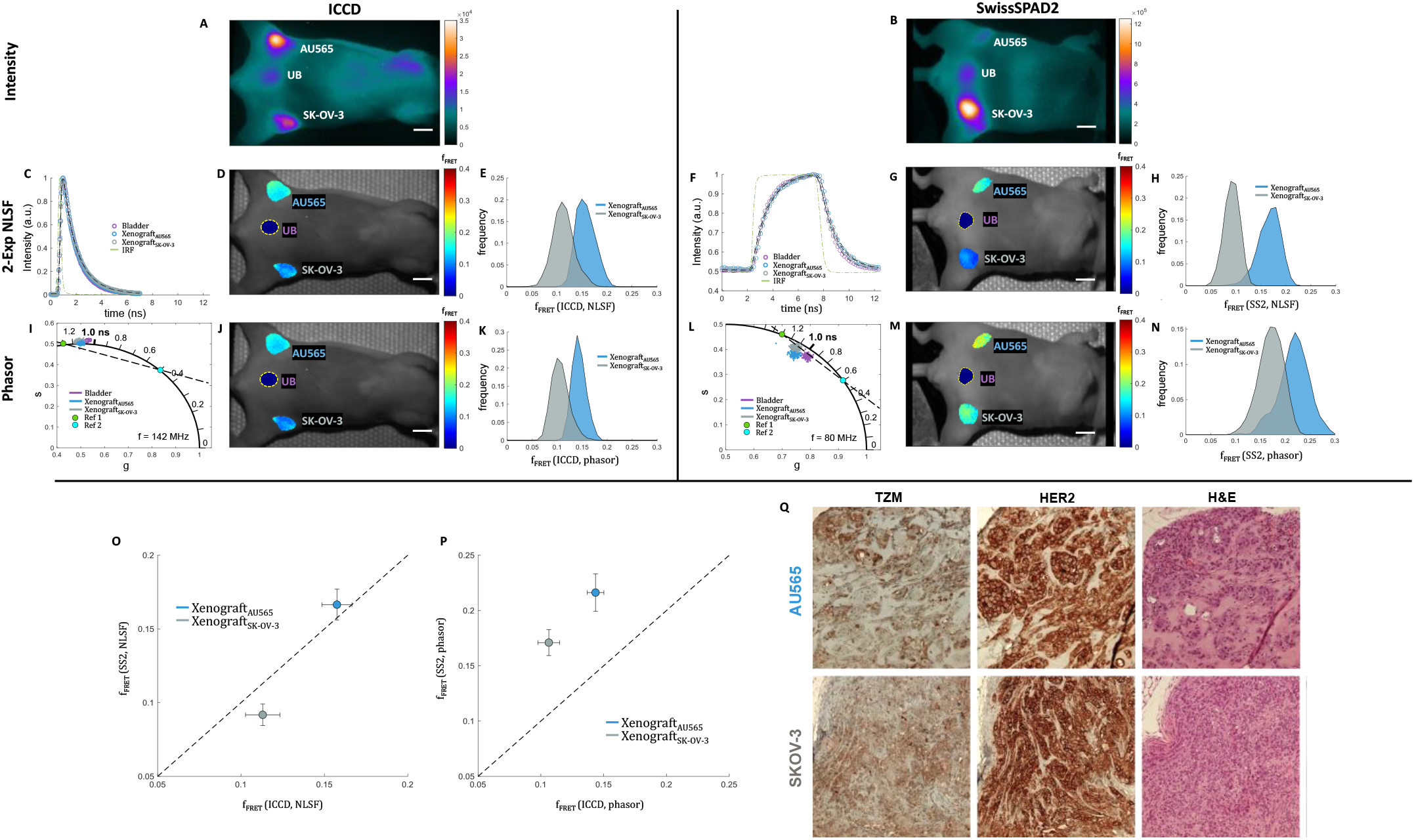
*in vivo* Trastuzumab-Her2 receptor engagement. Mice were injected with 20 µg of AF700-TZM and 40 µg of AF750-TZM and imaged by MFLI at 24 h post-injection (p. i.). Mouse 1 was imaged with the ICCD and mouse 2 with SS2. A, B: Fluorescence intensity images. A: ICCD; MCP voltage = 500V, integration time = 500 ms, illumination power = 2.13 mW/cm^2^. B: SS2; integration time = 2.65 s, illumination power = 3.2 mW/cm^2^;. C, F: ICCD and SS2 normalized whole ROI decays for the different organs, plotted with the corresponding IRF. D, G: MFLI FRET-fraction maps obtained by bi-exponential NLSF for both cameras. The urinary bladder (yellow dashed outline) was analyzed by 1-Exp NLSF and is therefore not included. E, H: Corresponding *f*_*FRET*_ KDE distributions for each well obtained for both cameras. I, L: phasor scatter plots color-coded by ROI, with overlaid reference lifetimes (green dot, reference 1; blue dot, reference 2) and dashed black line connecting them. J, M: pixel-wise phasor ratio maps and K, N: phasor-ratio KDE distributions for the two xenografts. O: Scatter plot (mean ± standard deviation) showing the FRET-fraction measured for each tumor with SS2 (mouse 2) versus that measured with the ICCD (mouse 1) retrieved through bi-exponential NLSF. P: Scatter plot of phasor ratio results (mean ± standard deviation) for ICCD and SS2. Scale bar in A, B, D, G, J, M: 6 mm. Q: *ex vivo* IHC of intracellular accumulation of TZM in AU565 and SK-OV-3 tumors in mouse 2. Consecutive sections were processed for H&E (showing cell localization and context), anti-HER2, and anti-TZM immunohistochemical staining. NovaRED was used as peroxidase substrate (brown stain), tissue was counterstained with methyl green. Scale bar = 100 µm.

For FRET quantification, bi-exponential NLSF analysis of the decays was performed in similar fashion to that of *in vitro* before (Fig. 2). Donor-only and quenched-donor lifetimes were retrieved through unconstrained full ROI decay fitting (*τ*_*1*_ = 1.3 ns *τ*_*2*_ = 0.5 ns). FRET fraction results obtained for mouse 1 (ICCD: Fig. 3D,E) and mouse 2 (SS2: Fig. 3G,H) are in high agreement. A direct comparison (Fig. 3O) shows that the two tumors behave remarkably similarly in both mice, with the AU565 xenograft exhibiting close to 50% more FRET (15.9 ± 1. 9% vs 10.4 ± 2.1%) than the SK-OV-3 (ovarian cancer) xenograft (Fig. S6 and Supplementary Note 2). Importantly, this pattern is preserved over time as shown in Figs. S8-S9, which summarize the analysis of mouse 2, observed at 48 hr p.i. (with SS2), and at 51 hr p.i (with the ICCD).

### Phasor Analysis

As *in vitro*, phasor ratio analysis (Fig. 3I-N) was performed using the reference lifetimes obtained in the NLSF analysis (*τ*_*1*_ = 1.3 ns *τ*_*2*_ = 0.5 ns). Noticeably, phasor ratio maps (Fig. 3J,M) and corresponding distributions for each tumor (Fig. 3K,N) demonstrate a remarkable consistency between methods and across mice.

### Immunohistochemistry

IHC analysis of excised tumors (Fig. 3Q) is in general agreement with MFLI data, showing more intracellular TZM accumulation in the AU565 tumors than in SK-OV-3 (Fig. 3Q left) despite high HER2 expression in both tumors (Fig. 3Q center). Reduced TZM accumulation in SK-OV-3 vs. AU565 tumors is consistent with decreased HER2-TZM binding as indicated by the lower FRET signal observed in SK-OV-3 vs. AU565 tumors via MFLI imaging with ICCD or SS2 detectors.

In summary, our experiments confirm our previous observation of a significant level of TZM-HER2 engagement in AU565 (breast cancer) tumor xenografts^43^. Moreover, we show that MFLI imaging can discriminate between different types of HER2 tumor models (breast AU565 and ovarian SK-OV-3) exhibiting distinct levels of TZM tumor delivery and HER2 binding, despite both tumor models showing HER2 overexpression. Our results suggest that these tumor models may display distinct tumor microenvironment challenges regarding effective drug delivery^47^. A manuscript focusing on the biological mechanisms underlying these differences in drug delivery between HER2 positive tumor models is currently in preparation.

### *In vivo* NIR MFLI-FRET measurements: Cetuximab-EGFR engagement in mouse xenografts

Cetuximab (CTM) is another monoclonal antibody used as EGFR inhibitor in clinical treatment of breast, colon, and head- and-neck cancers. As for TZM, AF700- and AF750-labeled CTM can be used in a NIR FLI-FRET assay to monitor CTM-EGFR drug-receptor engagement, as we recently demonstrated *in vitro*^48^. Here, similarly to the TZM experiment reported above, noninvasive MFLI-FRET imaging of nude mice bearing AU565 and SK-OV-3 tumor xenografts was performed. As described in *Online Methods*, two different mice were imaged at 48 hr p.i., either with the ICCD (mouse 1) or with SS2 (mouse 2). Fig. 4A,B shows the observed fluorescence intensity maps. In contrast to the TZM fluorescence experiment (Fig. 3), four regions are clearly visible: the liver as well as both tumor xenografts and the urinary bladder.

**Fig. 4:**
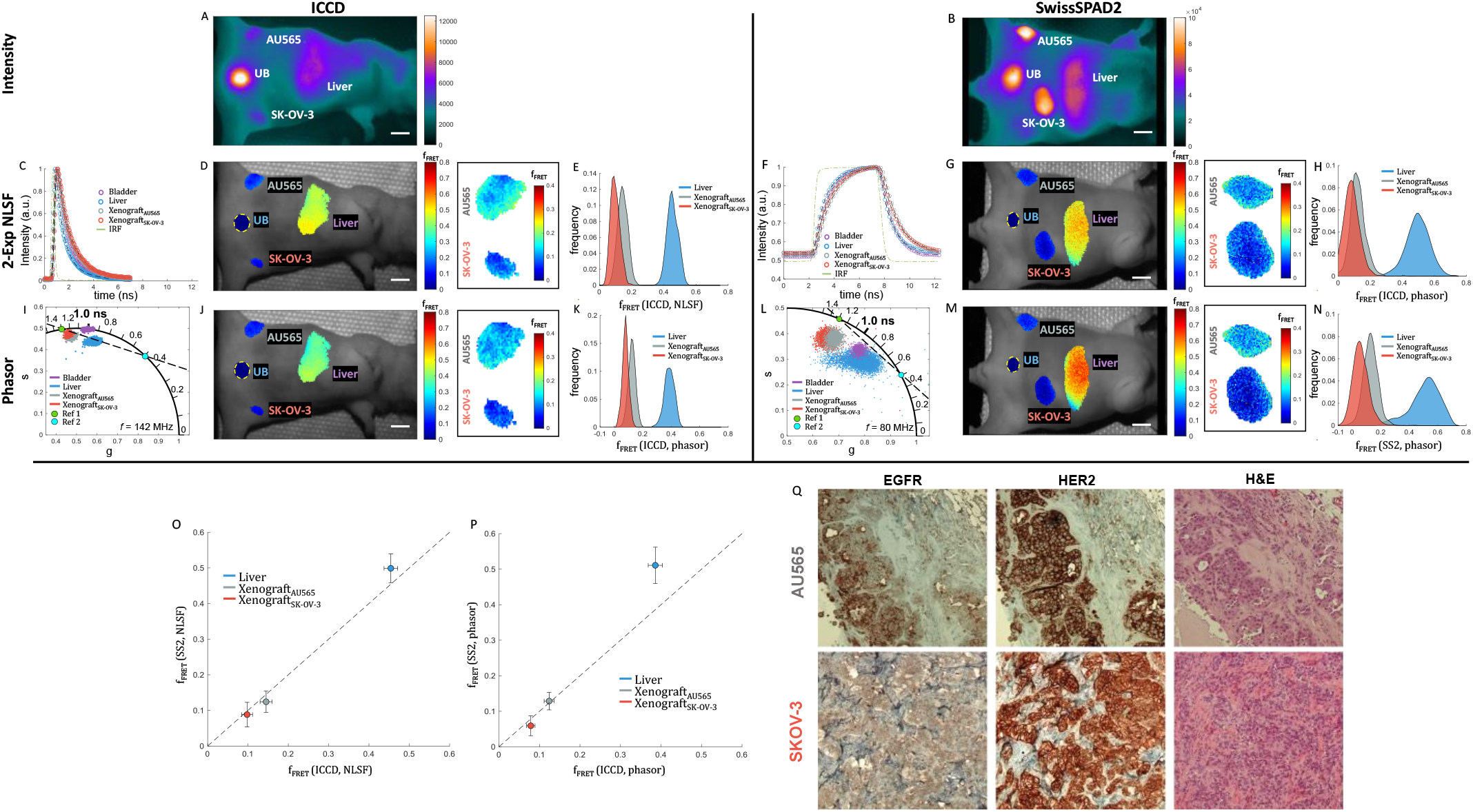
*in vivo* Cetuximab-EGFR engagement. Mice were injected with 20 µg AF700-CTM and 40 µg AF750-CTM and subjected to MFLI imaging at 48 h p.i. (mouse 1: ICCD, mouse 2: SS2). A, B: Whole body fluorescence intensity images for ICCD and SS2. A: ICCD; MCP voltage = 520 V, integration time = 500 ms, illumination power = 2.13 mW/cm^2^. B: SS2; integration time = 2.45 s, illumination power = 3.2 mW/cm^2^. C, F: ICCD and SS2 normalized whole ROI decays for the different organs, plotted with the corresponding IRF. D, G: MFLI FRET-fraction maps obtained by bi-exponential NLSF for both cameras. The urinary bladder (yellow dashed outline) was analyzed by 1-Exp NLSF and is therefore included as a constant 0 fraction. A zoomed in view of f_FRET_ quantification retrieved for both xenografts with color scale adjusted to match that used in Fig. 3 is shown on the right. E, H: Corresponding *f*_*FRET*_ KDE distributions for each well obtained for both cameras. I, L: phasor scatter plots color-coded by ROI, with overlaid reference lifetimes (green dot, reference 1; blue dot, reference 2) and dashed black line connecting them. J, M: Pixel-wise phasor ratio maps. A zoomed in view of phasor ratio quantification retrieved for both xenografts with color scale adjusted to match that used in Fig. 3 is shown on the right. K, N: Phasor-ratio KDE distributions for the two xenografts and the liver. O: Scatter plot (mean ± standard deviation) showing the FRET-fraction measured for each tumor and the liver with SS2 (mouse 2) versus that measured with the ICCD (mouse 1). P: Scatter plot of phasor ratio results (mean ± standard deviation) for ICCD and SS2. Scale bar in A, B, D, G, J, M: 6 mm. Q: *ex vivo* IHC validation of intracellular localization of EGFR and HER2 in AU565 and SK-OV-3 tumors in mouse 2. Consecutive sections were processed for H&E (showing cell localization and context), anti-HER2, and anti-EGFR immunohistochemical staining. NovaRED was used as peroxidase substrate (brown stain), tissue was counterstained with methyl green. Scale bar = 100 µm.

### NLSF Analysis

Following the workflow outlined in the *in vivo* TZM-HER2 experiment (Fig. 3), bi-exponential NLSF analysis of pixel-wise decays was performed using fixed component lifetimes obtained from whole ROI decay analysis (Fig. 4C,F: *τ*_*1*_ = 1.3 ns *τ*_*2*_ = 0.5 ns). As observed in the case of TZM FRET pair, the resulting FRET fraction quantification (Fig. 4D,G,E,H) in the SK-OV-3 tumor (*f*_*SK-OV-3*_ ∼ 8.9 ± 4.7 %) are smaller than those measured in the AU565 tumors (*f*_*AU565*_ ∼ 13.5 ± 4.4 %) and significantly lower than what is observed in the liver of both mice (*f*_*Liver*_ ∼ 48.5 ± 7.3 %), irrespective of the camera used. Comparison of results obtained in both mice using two different cameras (Fig. 4O) show once again exceptional agreement.

### Phasor Analysis

Pixel-wise phasor ratio analysis for each mouse (Fig. 4I,-N) shows a similar pattern to that observed previously for the TZM-HER2 experiment (Fig. 3-NL), with the notable difference of the presence of a liver phasor cluster. It is clear from both the phasor clusters themselves, as well as from the superimposed references (green dot, reference 1; blue dot, reference 2) anddashed black line connecting them, that tumors and liver phasors can be interpreted as a linear combination of a short (blue dot) and a long (green dot) single-exponential component.

Comparison of the FRET fractions obtained by this method with those obtained by NLSF analysis shows a truly remarkable correspondence, both between methods (compare Fig. 4D,E to 4J,K and 4G,H to 4M,N) and across mice (compare Fig. 4D,E to 4G,H and 4J,K to 4M,N). While there still exists a small offset between results obtained by the two methods, as indicated in Fig. 4O-P, the difference between tumors, and between tumor and liver is crucially preserved, further validating our approaches.

### Immunohistochemistry

The differences between tumors observed by MFLI-FRET are supported by IHC analysis (Fig. 4Q), which looked at anti-EGFR (left) and anti-HER2 (center) immunostaining, in addition to standard H&E staining (right). In contrast to the HER-TZM experiment, in which we monitored both the HER2 receptor and its respective antibody (TZM), here we examine only the receptors (EGFR and HER2) and not the actual probe (CTM). Therefore, we cannot directly confirm the uptake of CTM by AU565 or SK-OV-3 tumors. However, the comparatively low anti-EGFR staining in the SK-OV-3 tumor is consistent with the relatively lower FRET signal observed in SK-OV-3 xenografts.

In summary, our experiments confirm our previous observation of a significant level of CTM-HER2 engagement in AU565 (breast cancer) tumor xenografts^48^. The detection of a strong CTM FRET signal in the liver is also consistent with the known EGFR expression in mice livers. MFLI imaging with both ICCD and SS2 can therefore clearly discriminate different levels of CTM-EGFR drug-receptor tumor binding in two distinct EGFR-positive tumor models.

## III. Discussion

MFLI provides a unique contrast mechanism that enables monitoring key biological parameters dynamically and noninvasively. To perform such measurements in live specimen requires using NIR probes characterized by very short lifetimes, a challenge for current time-resolved detectors, which have lower detection efficiency in the NIR spectral range. In the past, time-gated ICCDs have been the workhorse for this type of MFLI studies. However, their high cost and limitations justify the exploration of alternative technologies. New development in SPAD technology has led to the integration of large time-gated CMOS SPAD cameras such as SwissSPAD2. These new cameras represent the next generation of imager for FLI studies across numerous biomedical applications. Using such cameras in the NIR spectral range is a priori challenging, and compounded by additional specificities of SS2 such as gate duration or rise- and fall-times, which are longer or of the order of magnitude of the lifetimes to be measured. We show here that these are not obstacles to accurate lifetime measurements. Indeed, the gate step size and the photon count, more than the gate duration or its rise- and fall-times, are the primary parameters affecting the measured lifetime precision^18,24,25^.

SS2 recovered FLI parameters of interest with high accuracy and in remarkable agreement with a state-of-the-art gated-ICCD. The outstanding performance of SS2 for MFLI applications ranges from resolving lifetimes of only a few hundreds of picosecond and lifetime differences of a few tens of ps. More importantly, this performance extends to multi-exponential decays analysis, where SS2 provides accurate estimates of FRET donor fraction.

These observations are further supported by *in vivo* studies which provided quantitative measurements of FRET donor fraction, a direct readout of drug-target engagement, over the whole animal body. SS2-measured FRET fractions were consistent with the molecular biology expected in the different organs in which signal was detected, including tumor xenografts with various target expression levels. Importantly, all *in viv*o experiments were performed at low illumination power (∼2.5 mW/cm^2^), well beyond the Maximum Permissible Exposure (MPE) limit, and with excellent SNR (∼400-1200, see Supplementary Tables 2-5)–even in deep seated tissues. This suggests that faster imaging rates can be achieved for applications with less demanding SNR constraints, potentially achieving live imaging capabilities.

By contrast with ICCDs, SS2’s technology offers room for significant future improvements: micro lenses can be added for up to 4-fold increased collection efficiency^24^, and upcoming next-generation SwissSPAD detectors will feature (i) shorter gate duration for better resolution, (ii) dual-gate architecture for more efficient photon collection and (iii) rolling-shutter recording, as well as (iv) higher recording rate to match that of the excitation laser, resulting in higher duty cycle. Moreover, the built-in field-programmable gate-array used to control the detector can be used for additional data pre-processing, opening the door for additional gains in speed and performance.

Our work presents a systematic comparison of two improved MFLI analysis approaches: (i) single- and bi-exponential NLSF analysis using full-period periodic convolution using a measured IRF and global lifetime constraints and (ii) phasor analysis based on calibration using a measured IRF. We demonstrate that these improvements capture the effects of sample topography and enable robust phasor analysis of target-receptor engagement *in vivo*.

In summary, we have validated a new large-area time-resolved SPAD camera for *in vitro* and *in vivo* macroscopic fluorescence lifetime imaging, supported by two complementary analysis approaches. This comprehensive investigation establishes the quantitative performances of SwissSPAD2 for NIR FLI imaging in complex scenarios. Due to its excellent performance, room for significant improvements, small form factor footprint and reduced cost, SwissSPAD2 and the next generation of time-gated detectors are expected to become the technology of choice for macroscopic FLI imaging with impact on a wide-range of applications – ranging from preclinical studies in drug development to optical guided surgery.

## IV. Online Methods

### Macroscopic FLI with SwissSPAD2

An illustration of the SwissSPAD2 MFLI configuration (reflectance geometry) used herein is provided in Fig. 5. Technical details about the camera can be found in earlier publications^22–24^ and are briefly summarized below. The system’s excitation source was a tunable Ti-Sapphire laser (Mai Tai HP, Spectra-Physics, CA, USA). Laser excitation was directed to the sample plane directly from a multimode optical fiber (QP200-2-VIS-NIR, Ocean Optics, FL, USA) output. Emitted fluorescence was collected through an application-specific bandpass emission filter (Supplementary Fig. S10) by a macroscopic photographic lens (AF Nikkor 50mm f/1.8D, Nikon, Tokyo, Japan). IRFs (instrument response functions) were acquired similarly, using a sheet of white paper as sample (for *in vitro* experiments) or the mouse itself (in the case of *in vivo* measurements), after removing the emission filter. Slightly lower laser power and integration time were also used. For use with SS2, which requires synchronization with a source signal at around 20 MHz, a frequency divider (TOMBAK, Laser Lab Source Corporation, MT, USA) was used to divide the Mai Tai’s laser repetition rate (∼80 MHz) by a factor of four (*f*_*SYNC*_ = 19.77 MHz reported). SS2 was set to acquire 10-bit gate images consisting of 1,020 accumulated 1-bit gate images, each 1-bit image resulting from exposure of each SPAD pixel to the incoming photon flux for a user-specified duration of *n*_*E*_ × 400 ns, where *n*_*E*_ is some number typically between 1 and 100^23^. During that 1-bit accumulation period, each SPAD is “on” (*i*.*e*. capable of detecting a photon) for a duration *W*_*SS2*_ (gate width) and “off” the remainder of each *T*_*sync*_ = 1/ *f*_*SYNC*_ = 50.6 ns period, and able to detect at most one photon. Each full gate image differs from the previous one by its distance to the laser pulse, or gate offset, by a user-specified amount *dt* (“gate step”), multiple of 1/56 ns = 17.857 ps. *δt* = 178.57 ps was used in this work. Although gate images covering the whole 50 ns window of SwissSPAD2 max sync rate (*i*.*e*. 280 gates in total) were acquired, resulting in the acquisition of four laser periods-worth of data, a single laser period-worth of data (70 first gates) was used in most analyses, unless mentioned otherwise. Because the laser period *T* = 12.65 ns is comparable to the shortest gate implemented in SS2 (*W*_*SS2*_ = 10.7 ns), which results in severely deformed periodic decays, a larger gate duration (*W*_*SS2*_ = 17.9 ns) was chosen to minimize this artifact (Supplementary Note 3 & Supplementary Fig. S11). To increase the total effective integration time for each gate, each acquisition was repeated multiple times (50-100 repetitions). Integration time for each gate image is calculated following ref.^23^ as: *T*_int_ = (8*n*_*G*_ −1)*bNT*_*Sync*_, where *n*_*G*_ is a gate sequence parameter (typically *n*_*G*_ = 100), *b* is the number of 1-bit gate images per final image (*b* = 1,020), *N* is the number of accumulations (e. g. ***N*** = 70) and *T*_*sync*_ = 1/ *f*_*SYNC*_ is the period of the synchronization signal. *T*_*int*_ was set between 1.02-4.08 s depending on the sample brightness. Actual values of these parameters are reported when needed and are available in the raw data provided on Figshare^49^. Data is transferred asynchronously via a USB 3.0 connection from the SS2’s field-programmable gate-array (FPGA) to the computer under control of a dedicated open-source LabVIEW program, [SwissSPAD Live](https://swissspad-live-distribution.readthedocs.io/en/latest/)^50^.

**Fig. 5:**
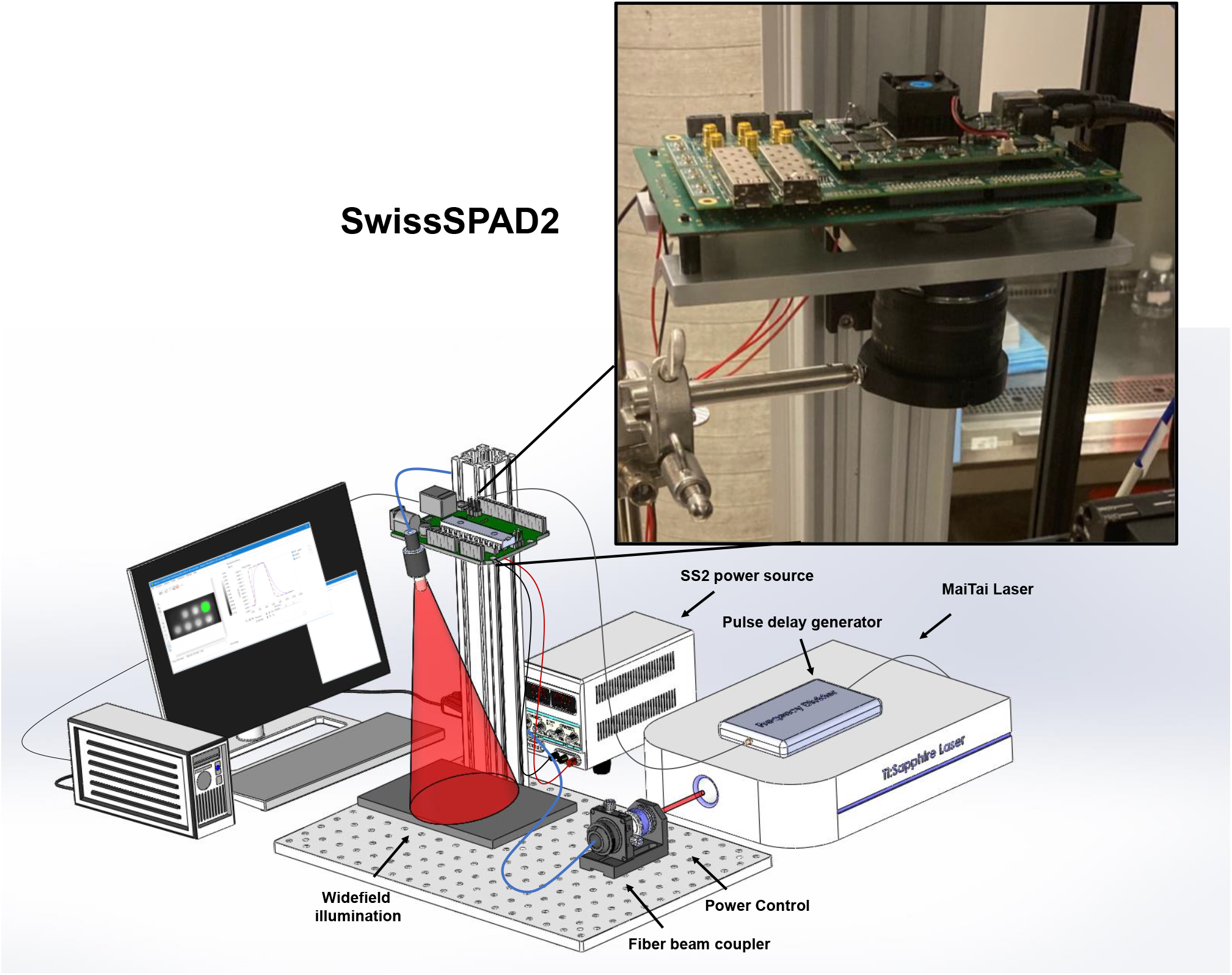
Schematic illustration of the widefield time-resolved imaging system equipped with the SwissSPAD2 camera. All imaging was performed in reflectance geometry. A fs pulsed, tunable Ti:Sapphire laser beam was directed through a power control module before coupling into a multimode fiber (blue line). The divergent output of the fiber was directed toward the sample to achieve a Gaussian illumination profile. Light emitted by the sample was collected using an overhead macroscopic objective lens directly attached to the C-mount port in front of SwissSPAD2 (SS2, green PCB boards and photograph inset). SS2 is powered by two regulated external power supplies (red lines) and a small DC adapter. A TTL pulse derived from the laser trigger signal by a frequency divider module is used to synchronize data acquisition (thick black line). Data is transferred to a PC via a USB 3 cable (thin black cable).

### Macroscopic FLI via Gated-ICCD

MFLI was also performed using a widefield time-resolved FLI apparatus equipped with a gated-ICCD (described previously^51^). In brief, the system used the same excitation source as described previously, but a digital micro-mirror device (DLi 4110, Texas Instruments, TX, USA) was used for widefield illumination over the sample plane. The time-gated ICCD (Picostar HR, LaVision, GmbH, Bielefeld, Germany) was set to acquire gate images with a gate width of *W*_*ICCD*_ = 300 ps, separated gate steps *δt* = 40 ps (details provided elsewhere^52^). Data was acquired over a window of duration shorter than the full laser period (generally *G* = 176 total gate images per acquisition, *i*.*e. D* = 7 ns, but occasionally longer, see details in each figure caption). As with SS2, IRFs were acquired with equivalent conditions to those used for fluorescence imaging. During fluorescence imaging, fluorophore-dependent filters were inserted, the ICCD’s microchannel plate (MCP) voltage was increased for sufficient signal amplification (between 350-550 V depending on sample brightness) and integration time adjusted within the range 300-500 ms per gate image. Details are provided in each figure caption. Because no calibration of the camera gain (photon per camera unit signal) as a function of MCP voltage was performed for this device, no signal-to-noise ratio is reported.

### Well-plate sample preparation and imaging

- *AF750 serial dilution*. To test SS2’s capability to quantitatively resolve short-lifetime NIR dye at low concentrations, a serial dilution of AF750 in two distinct aqueous buffers (H_2_O and PBS) was prepared in a well-plate (Fig. S2). Concentrations ranged from 25 µg/mL to 3 µg/mL. Laser excitation was set to 750 nm and the emission filter used was 780 ± 10 nm (Semrock, FF01-780/12-25).
- *IRDye 800CW-2DG*. A well-plate sample of IRDye 800CW-2DG (LI-COR Biotechnology, NA, USA) was prepared at a constant concentration of 15 μM using seven different buffers: Intracellular pH Calibration Buffer Kit (pH 4.5, pH 5.5, pH 6.5, pH 7.5 – ThermoFisher, USA) distilled H_2_O, PBS (pH 7) and DMSO. Laser excitation was set to 760 nm and the emission filter used was 800 ± 10 nm (Semrock, FF01-800/12-25).
- *AF700/AF750 IgG FRET*. A well-plate was prepared using the FRET pair AF700/AF750 IgG (ThermoFisher Scientific, MA, USA) at four acceptor-donor ratios (0:1-3:1) in PBS buffer. Each well contained AF700 prepared at an equivalent concentration (30 µg/mL) across all wells. The well-plate was imaged ∼24 hours following preparation. Laser excitation was set to 700 nm and the emission filter used was 740 ± 10 nm (Semrock, FF01-740/13-25). Laser power was set to 1.75 mW for the SS2 and 0.8 mW for the ICCD.
- *TZM-AF700 and CTM-AF700 (donor only)*. A well-plate was prepared using AF700 conjugated to either TZM or CTM (Fig. S7). The concentration was kept constant for both TZM-AF700 and CTM-AF700 (33 µg/mL) across separate rows. Buffer pH was varied from 4.5 to 7.5 as explained previously for the IRDye 800CW-2DG well-plate experiment. Laser excitation was set to 700 nm and the emission filter used was a 715 nm longpass filter (Semrock, FF01-715/LP). Laser power was set to 1.2 mW.

### Animal experiments

All animal procedures were conducted with the approval of the Institutional Animal Care and Use Committee at both Albany Medical College and Rensselaer Polytechnic Institute. Animal facilities of both institutions have been accredited by the American Association for Accreditation for Laboratory Animals Care International. HER2-overexpressing cell lines AU565 and SK-OV-3 used in this study were obtained from ATCC (Manassas, VA, USA) and cultured in RPMI and McCoy’s media respectively supplemented with 10% fetal bovine serum (ATCC) and 50 Units/mL/50 μg/mL penicillin/streptomycin (Thermo Fisher Scientific, Waltham, MA, USA). Tumor xenografts were generated by injecting 10×10^6^ AU565 cells and 4×10^6^ SK-OV-3 in phosphate-buffered saline (PBS) mixed 1:1 with Cultrex BME (R&D Systems Inc, Minneapolis, MN, USA) on the opposite sides of inguinal mammary fat pads of female 5-week-old athymic nude mice (CrTac:NCr-Foxn1nu, Taconic Biosciences, Rensselaer, NY, USA). Tumors were monitored daily over a period of 3-4 weeks.

Two xenograft-bearing mice were injected with 20 µg AF700-TZM and 40 µg AF750-TZM. Each mouse was imaged separately using either the SS2 or the ICCD at 24 h post-injection (Fig. 3, ∼ 1hr between). Additionally, one of these mice was imaged at 48 h (SS2) and 51 h (gated-ICCD) post-injection (i.e., three hours after SS2 imaging, Supplementary Fig. S8&S9). Another two xenograft-bearing mice were injected with 20 µg AF700-CTM and 40 µg AF750-CTM. Each mouse was imaged separately using either the SS2 or the ICCD at 48 h post-injection (Fig. 4, ∼ 1 hr between session). All injections were performed retro-orbitally on anesthetized mice. During mouse imaging, isoflurane anesthesia was performed, and the body temperature of each animal was maintained using a warming pad (Rodent Warmer X2, Stoelting, IL, USA) on the imaging plane.

### Immunohistochemical analysis

Excised tumors were fixed in formalin, paraffin embedded, and processed for IHC. Epitope retrieval was performed by boiling deparaffinized and rehydrated 5µm sections in 1 mM EDTA pH 8.0 for 30 min. IHC staining was carried out using a standard protocol from Vectastain ABC Elite kit (Vector Labs, Burlingame, CA, P/N: PK-6101). Vector NovaRED (Vector Labs) was used as a peroxidase substrate. Tissue sections were counterstained with Methyl Green (Sigma, P/N: M8884). Hematoxylin Eosin stain was used for basic histology. Primary antibodies were as followed: rabbit monoclonal HER2 1:800 (Cell Signaling, P/N: 2165), rabbit monoclonal EGFR 1:50 (Cell Signaling, P/N: D38B1), rabbit monoclonal TZM 1:100 (R&D Systems, P/N: MAB95471-100). Brightfield images were acquired using Olympus BX40 microscope equipped with Infinity 3 camera (Lumenera Inc., Ottawa, ON, Canada).

### Preprocessing of SS2 data

All SS2 data was processed using [Alligator](https://sites.google.com/a/g.ucla.edu/alligator/)^18^ software as described in Supplementary Note 4. Briefly, raw gate images (consisting in the sum of four 8-bit accumulation of individual 1-bit frames) were corrected for pile-up effects as described previously^24^. Next, subtraction of a detector background files acquired similarly as the data were used to correct for detector background (dark count noise) inhomogeneities and minimize the influence of “hot” SPADs/pixels characterized by intrinsically high dark count noise (Supplementary Fig. S12). Only one laser period (12.5 ns) worth of data was retained from each dataset (first 12.5 ns i.e. first 70 gates) except for one experiment (CTM *in vivo*). For the CTM *in vivo* experiment, significant photobleaching was observed during the last half of imaging. Hence, for this experiment’s data processing, three 12.5 ns sections (first 70 gates, 71-140 gates and 141-210 gates) of MFLI data were extracted from the sum of the first twenty acquisitions. These three data were summed together to provide 60 total accumulations. 2×2 binning was used for all SS2 data herein. IRF data was processed in the same manner as fluorescence data (pile-up correction and background correction). SNR numbers reported in the text correspond to the square-root of the resulting pile-up and background-corrected pixel values.

### Preprocessing of gated-ICCD data

All ICCD data was binned spatially (using either 2×2 or 4×4 binning). No further preprocessing was undertaken for phasor analysis. For NLSF analysis, because the decays were truncated to cover only 7 ns out of the 12.5 ns of a full laser period, the IRFs of individual pixels were extrapolated to cover the full-period (see Supplementary Note 4 for details). No further preprocessing of fluorescence decays was performed.

### Nonlinear Least Square Fit of Fluorescence Decays

All full period (SS2) or partial (ICCD) decays {*G*_*i*_}_*i* = 1,..,*N*_ were fitted using the Levenberg-Marquardt non-linear least square fit (NLSF) algorithm implemented in [Alligator](https://sites.google.com/a/g.ucla.edu/alligator/)^18^. Briefly, experimentally acquired pixel-wise IRF data (SS2) or extrapolated *T*-periodic IRF data (ICCD, see Supplementary Fig. S5), *I*_*T*_ (*t*), was utilized for *T*-periodic convolution with a single- or bi-exponential *T*-periodic decay model *F*_*T*_(*t*), including an optional IRF offset parameter *t*_*0*_ (set to 0 in all our analyses) and a baseline parameter *B* accounting for residual uncorrelated background (Eq. (1)) (Supplementary Note 4):

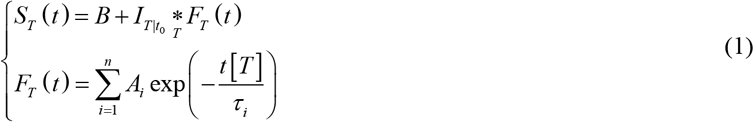

where the notations of ref.^46^ are used. In particular, 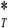 denotes the cyclic convolution product, 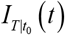 designates the experimental *T*-periodic IRF (with unknown offset *t*_*0*_) and *x*[*T*] denotes *x* modulo *T*. Index *T* in the function notation indicates a *T*-periodic function. *A*_*i*_ and *τ*_*i*_ are the amplitude and lifetime of component *i*, where the number of components *n* is either 1 or 2 depending on the experiment. Weighted fits were performed using the minimization function (Eq. (2)):

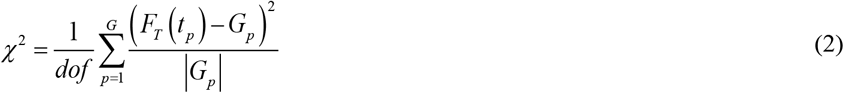

where *dof* is the number of free parameters of the fit, *t*_*p*_ is the *p*^th^ gate location in the period and *G*_*p*_ the gate value. If *G*_*p*_ = 0, a weight of 1 replaces the factor | *G*_*p*_ | in Eq. (2).

Note that in the *in vivo* case, in which the mouse itself was used for IRF acquisition, the corresponding recorded decay is not expected to be the true IRF, as the collected signal corresponds mostly to photons scattered off the mouse surface (or the most superficial layers of the skin). It does not account for propagation and scattering of the excitation light deep into tissues, nor does it account for propagation and scattering of the emitted fluorescence through the same layers of tissue^53^. Nevertheless, as discussed in Supplementary Note 5 and illustrated in Supplementary Fig. S13, which represents the local delay of the recorded scattered laser signal (relative to that measured with a white sheet of paper) as an equivalent topographic map (see below), the corresponding signal does differ significantly from a mere “paper IRF” and is therefore expected to be a better approximation of the true IRF than a simple paper IRF.

### FRET fraction analysis using decay fits

As discussed in ref.^46^, a periodic decay *F*_*T*_(*t*) expressed according to Eq. (1), is the infinite summation of the non-periodic decay:

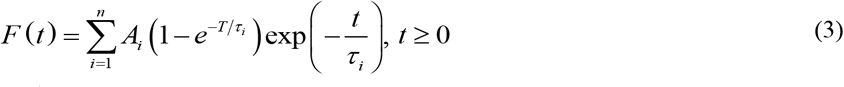

whose component *i* contributes 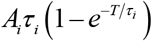 to the total integral over [0, +∞] and whose intensity fraction *f*_*i*_ is therefore:

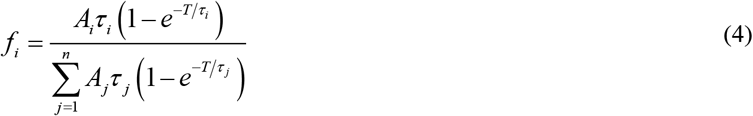

This intensity fraction can be directly compared to that obtained by phasor ratio analysis (Eq. (8) below).

### Phasor Analysis

All phasor analysis was performed using [Alligator](https://sites.google.com/a/g.ucla.edu/alligator/)^18^ as detailed elsewhere^23,34^. In brief, for every pixel with coordinate (*x, y*) within the MFLI region of interest, the uncalibrated *discrete* phasor *z*(*x, y*) = *g*(*x, y*) + *i s*(*x, y*), where *i* the complex root of −1, was retrieved from the time-gated decay {*G*_*p*_(*x, y*)} _*p* = 1,..,*G*_ according to Eq. (5).

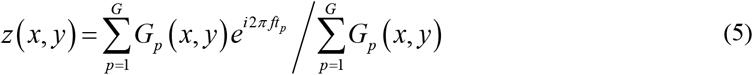

where *f* is the phasor frequency *G* the number of gates, *G*_*p*_ the *p*^th^ gate value at pixel (*x, y*). In particular, phasors computed for ICCD data, where truncated decays were recorded, correspond to the truncated (and offset) decay case discussed in ref.^46^. The harmonic frequency *f* was chosen equal to 1/*D* ∼ 142 MHz – where D is the time span of the recorded decay. The resulting appearance of the phasor plot differs from the standard case, where the phasors of single-exponential decays are located on the so-called universal semicircle (or universal circle, UC)^45^. Instead, in the case of a truncated decay, the single-exponential decay’s phasor locus (or SEPL^34^) is in general a complex curve, which, after calibration, may or may not overlap partially with the UC (see next).

For SS2, one laser period-worth of data was used, therefore *D* = *T*, and *f* = 80 MHz. However, since the number of gates (*G* = 70) is finite, the SEPL is also expected to depart from the UC and be closer to an arc of circle with larger radius than the UC^45^. In practice, *G* is sufficiently large for this difference to be negligible, and the UC was used in all representation of calibrated phasors.

### Phasor calibration

Visual depiction of how phasor calibration was undertaken is illustrated in the *Supplementary Material* (Supplementary Fig. S14 and S15 for SS2 and ICCD, respectively) and follows ref.^46^. Briefly, the uncalibrated phasor of the IRF data corresponding to the sample of interest (a sheet of white paper for the *in vitro* samples, and the mouse itself for *in vivo* measurements [Supplementary Fig. S16]) was calculated using Eq. (5) on a pixel-by-pixel basis. These *calibration* phasors, 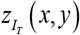, associated with lifetime *τ*_*IRF*_ = 0 were used to compute the calibrated phasor of sample data, 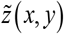, according to Eq. (6):

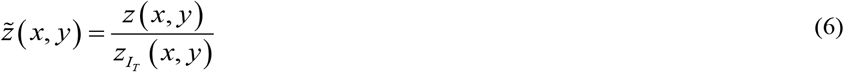

As discussed above, due to their discrete nature and the nontrivial shape of the IRFs involved in their calculation, calibrated phasors of single-exponential decays, normally expected to be located on the so-called universal circle (UC), are instead located on a slightly different curve^18^ (dubbed SEPL – or Single-Exponential Phasors Locus - in ref.^34^), which is expected to depart from the UC for large enough lifetimes. In practice, for the number of gates used and the short lifetimes studied in this work, the SEPL is indistinguishable from the UC, which was used instead.

### Phase lifetime calculation

The phase lifetime, *τ*_*ϕ*_, was calculated using the components (*g, s*) of the calibrated phasor (Eq. (6)) according to Eq. (7), which is only correct for an infinite number of gates *G*, but is a good approximation in the experimental situations described in this work^34^:

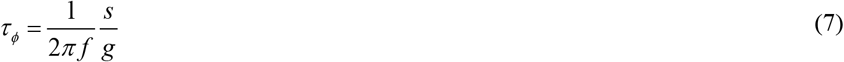

### Phasor ratio calculation

In the case of decays comprised of the contribution of two different species, such as encountered for the mixtures of quenched donor and unquenched donor lifetimes in the FRET assays presented here, the phasor of the mixture is a linear combination of the phasor of each species^18,45,46^. The intensity fraction of each species in the mixture can be graphically recovered from the location of the phasor with respect to the pure species (reference) phasors. In the case studied here, one of the reference phasor, the donor-only – or unquenched donor – phasor can be measured experimentally with the pure donor sample, while the second reference can be inferred from the observed *linear arrangement* of the phasors of the different acceptor to donor ratio samples (see Fig. 2): while there is no guarantee – and in fact it is unlikely – that the decay of a maximally quenched donor is characterized by a strictly single-exponential decay, a natural reference to use for quantification of the mixture is the intersection of the best-fit line on which all samples are aligned, with the SEPL (see Supplementary Fig. S17 for a graphical illustration). Since there are two intersections, that with the shortest lifetime is chosen as the second reference (the other intersection should correspond to the donor-only sample if it is characterized by a single-exponential decay – this is approximately verified here, but not exactly, which is why the actual donor phasor is used as the first reference). Because of shot noise and other sources of variance, each phasor is projected orthogonally onto the line connecting both references to extract the so-called phasor ratio *r* expressing the relative distance of the phasor to the two reference phasors (Eq. (8)):

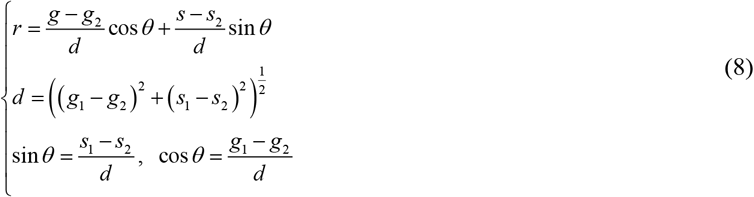

where (*s*_*i*_, *g*_*i*_), *i* = 1 or 2, correspond to the phasor of the two references and (*g, s*) is the mixture phasor. The phasor ratio *r* defined in Eq. (8) corresponds to the intensity fraction of species 1 in the mixture^18^ and can thus be compared to the intensity fraction extracted by NLSF of decays (Eq. (4)), assuming that each component species is characterized by a single-exponential decay.

### Statistical Analysis

Raw analysis data exported from AlliGator as text files were subsequently processed in MATLAB to obtain publication-quality plots and statistical representations. Boxplots used in Fig. 1 were created using MATLAB with all solid black lines marking the average value and each box width marking 1-standard deviation. Kernel density estimation distributions were calculated in MATLAB using the lower and upper bounds of the listed x-axis and a 10^−2^ interval (e.g., [0:0.01:0.8] for Fig. 4E) for all cases herein. All average and standard deviation results used for comparative scatter plots were calculated using MATLAB (dashed black line marks the diagonal in all cases). Linear regression used in Fig. 1Q&R (light blue dashed line) was performed in MATLAB. All average and standard deviation results were computed using MATLAB. Example script is provided for reproduction purposes^49^.

### Fluorescent probes

IRDye 800CW 2-DG was purchased at Li-Cor (Lincoln, Nebraska, USA). Cetuximab (CTM) was purchased at MedChemExpress (Monmouth Junction, NJ, USA), trastuzumab (TZM) was obtained through Albany Medical Center pharmacy. Fluorescent labeling of TZM is described in depth elsewhere^43^. Fluorescent labeling of CTM was performed following protocol described elsewhere^54^. Mouse IgG AF700 and rabbit IgG AF750 were purchased at Thermo Fisher Scientific (Waltham, MA, USA).

## V. Associated content

The supplementary information is available free of charge at https://XXXXXXX

Software and Data Availability: All analyses are performed using freely available software^18^, and data as well as analysis details and results are available on a public cloud repository in order to ensure reproducibility^49^.

## VII. Acknowledgments

This work was supported by the National Institute of Health Grants (R01CA237267, R01CA207725 and R01CA250636) and in part by the UCLA Jonsson Comprehensive Cancer Center Seed Grant Program. A.U. was supported through the Swiss National Science Foundation under grant 200021-166289.

We acknowledge the support of Albany Medical Center pharmacy for donating trastuzumab used herein. JTS would like to thank Ms. Marien Ochoa-Mendoza for valuable discussion and insight.

