## Supplementary Information for "*In vitro* and *in vivo* NIR Fluorescence Lifetime Imaging with a time-gated SPAD camera"

### Supplementary Tables

#### Supplementary Table 1: Lifetime values (mean ± standard deviation) corresponding to Fig. S2E, H, L, O.

| Conc (µg/mL) | 3 | 8 | 16 | 25 |
|---|---|---|---|---|
| $\tau_{SS2, NLSF}$ (ns) | 0.58 ± 0.06 | 0.66 ± 0.04 | 0.68 ± 0.03 | 0.68 ± 0.02 |
| $\tau_{SS2\ Phasor}$ (ns) | 0.56 ± 0.06 | 0.64 ± 0.04 | 0.65 ± 0.03 | 0.65 ± 0.03 |
| $\tau_{ICCD\ NLSF}$ (ns) | 0.58 ± 0.06 | 0.63 ± 0.04 | 0.65 ± 0.02 | 0.65 ± 0.02 |
| $\tau_{ICCD\ Phasor}$ (ns) | 0.56 ± 0.05 | 0.60 ± 0.04 | 0.62 ± 0.02 | 0.62 ± 0.02 |

#### Supplementary Table 2: SNR values (mean ± standard deviation) corresponding to Fig. S2.

| **Conc (µg/mL)** | **SNR** |
|---|---|
| 25 µg/mL$_{(H2O)}$ | 1853.01 ± 341.04 |
| 25 µg/mL$_{(PBS)}$ | 1741.14 ± 303.85 |
| 16 µg/mL$_{(H2O)}$ | 1907.07 ± 307.31 |
| 16 µg/mL$_{(PBS)}$ | 1803.82 ± 281.34 |
| 8 µg/mL$_{(H2O)}$ | 1510.36 ± 214.98 |
| 8 µg/mL$_{(PBS)}$ | 1519.94 ± 207.64 |
| 3 µg/mL$_{(H2O)}$ | 1226.84 ± 122.96 |
| 3 µg/mL$_{(PBS)}$ | 1210.41 ± 125.81 |

#### Supplementary Table 3: SNR values (mean ± standard deviation) corresponding to Fig. 1.

| **Well ROI** | **SNR** |
|---|---|
| DMSO | 1320.59 ± 81.03 |
| H2O | 1135.15 ± 85.47 |
| PBS | 1036.37 ± 79.65 |
| pH 4.5 | 1122.30 ± 96.33 |
| pH 5.5 | 1086.64 ± 73.71 |
| pH 6.5 | 1087.45 ± 68.47 |
| pH 7.5 | 1005.41 ± 78.46 |

#### Supplementary Table 4: SNR values (mean ± standard deviation) corresponding to Fig. 2.

| **Well (A:D)** | **SNR** |
|---|---|
| (0:1) | 844.24 ± 96.28 |
| (1:1) | 810.00 ± 80.10 |
| (2:1) | 719.73 ± 72.32 |
| (3:1) | 605.90 ± 52.08 |

#### Supplementary Table 5: SNR values (mean ± standard deviation) corresponding to Fig. 3, 4 and S8-9.

| Animal imaging session | Liver | Bladder | Xenograft$_{AU565}$ | Xenograft$_{SK\text{-}OV\text{-}3}$ |
|---|---|---|---|---|
| Fig. 3 – (Mouse$_{TZM}$ day#1) | N/A | 1237.67± 34.64 | 1040.58 ± 34.71 | 1735.14± 143.71 |
| Fig. 4 – (Mouse$_{CTM}$ day#2) | 492.70 ± 27.20 | 584.81 ± 32.17 | 531.68 ± 47.35 | 492.70 ± 27.20 |

| | | | | |
|---|---|---|---|---|
| Fig. S8-9 – ($Mouse_{TZM}$ day#2) | N/A | 540.34 ± 24.03 | 431.56 ± 29.68 | 607.76 ± 27.27 |

### Supplementary Figures

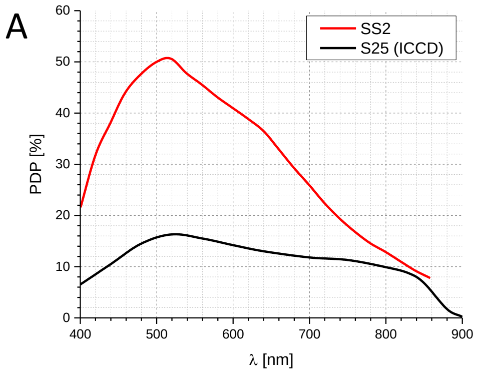

B

| | ICCD | SS2 |
|---|---|---|
| Fill Factor (FF) | ~1 | 0.105(*) |
| Gate Duration (ns) | 0.3 | 17.9 |
| Gate Period (ns) | 12.5 | 50 |
| Duty Cycle (DC = W/T) | 0.024 | 0.36 |
| FF × DC | 0.024 | 0.038 |
| FF × DC x PDP @ 720 nm (%) | 0.28 | 0.83 |

**Fig S1:** Characteristics of the two detectors used in this study. A. Sensitivity. The photon detection probability of SwissSPAD 2 assumes a 7 V excess bias voltage [1]. The quantum efficiency of the S25 (aka SuperGen 2) photocathode manufactured by Photonis for LaVision is assumed to be identical to that of a similar photocathode developed by Photonis for a different type of detector [2]. B. Spatio-temporal factors affecting the photon detection efficiency of the two detectors. Spatial effect: fill-factor; Temporal effect: duty cycle (gate duration *W* divided by gate period *T*). For the ICCD, the gate repetition rate was identical to the laser period $T$ = 12.5 ns, while the gate width was $W$ = 300 ps. For SS2, the repetition rate was 4 times the laser period, or 50 ns, but the gate width was $W$ = 17.9 ns. The corresponding duty cycle is therefore larger for SS2. The product of both effects (FF × DC) is similar for both detectors. Overall, there is a ~ 3-fold advantage for SS2 at 720 nm when the photon detection probability is considered. (*) While the camera used in this work was not equipped with one, SS2 can be equipped with a microlens array, further enhancing SS2's photon detection efficiency by a factor close to 3 [3].

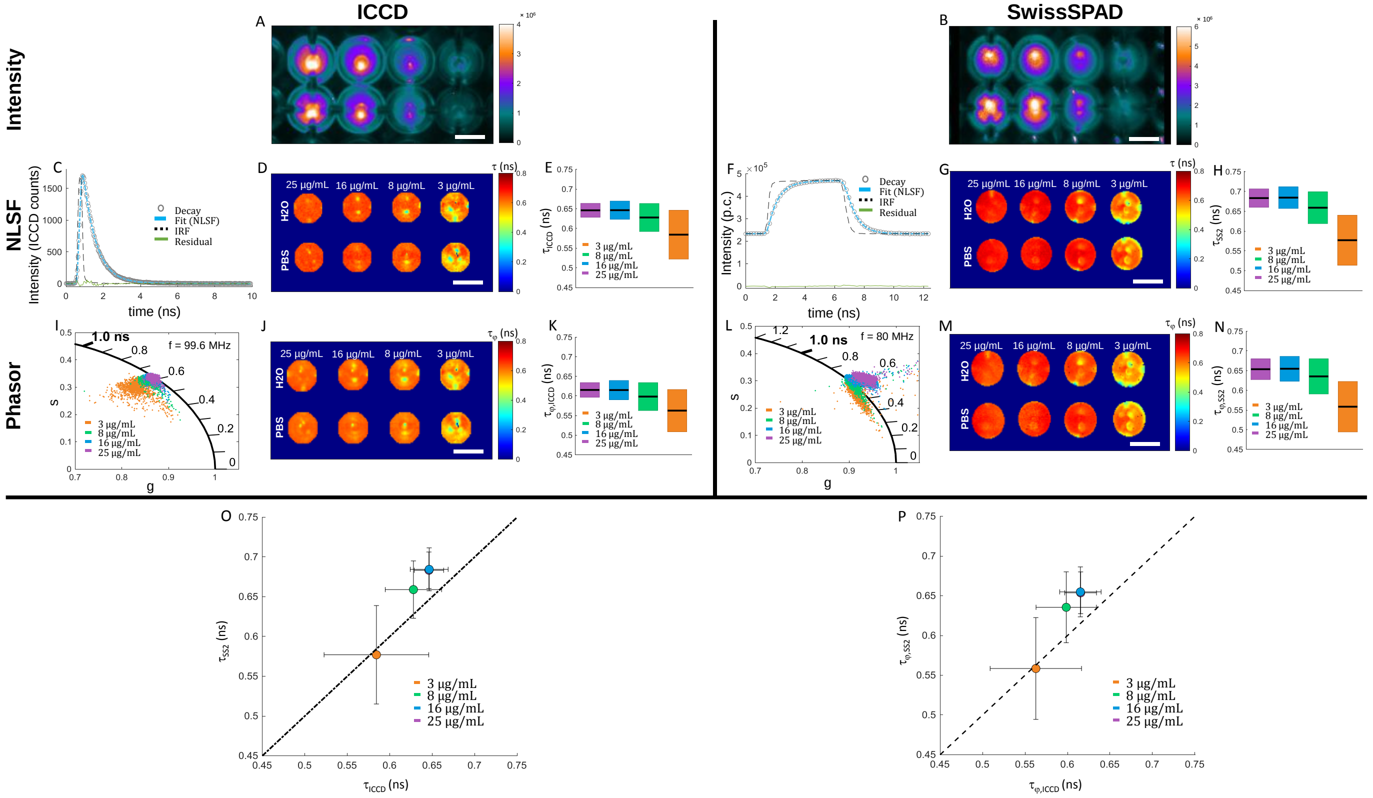

**Fig S2**. Serial dilution of AF750. A, B: Fluorescence intensity images; ICCD: MCP voltage = 440 V, integration time/gate image = 409 ms, illumination power = 0.95 mW/cm$^2$. SS2: integration time = 4.08 s, illumination power = 2.0 mW/cm$^2$. C, F: Representative IRFs (dotted black curves), decays (open circles), mono-exponential NLSF fits (dashed blue curves) and residuals (green curves). Note in particular the large SS2 gate, which results in a vertical offset of the decay due to a full period integration (*Methods*) and a characteristics "mirroring" of the fluorescence decay on the rising edge side. ICCD data (color scale in A, vertical axis in C) are represented in "CCD counts", which are proportional to the number of detected photons, while SS2 data (color scale in B, vertical axis in F) are directly measured in photon counts. D, G: lifetime maps obtained by NLSF. Spots and features visible in the intensity images as well as in the lifetime maps correspond to caustics formed by the wells' curved walls (see main text). E, H: Boxplots of the fitted lifetimes for all well concentrations (PBS and water-dissolved dye samples). The average value is indicated by a black segment, the standard deviation is equal to half the box height. Each well data is color coded as indicated in the legend. The larger dispersion of the fitted lifetimes at lower concentrations is due to shot noise effects. The negative bias at lower concentrations is due to the increasing influence of the lower lifetime hotspots in the distribution. I, L: IRF-calibrated phasors for all four AF750 concentrations. J, M: Phase lifetime maps obtained from I, L. K, N: Boxplot representing phase lifetime results for all well concentrations (PBS and water-dissolved dye samples). The average value is indicated by a black segment, the standard deviation is equal to half the box height. Each well data is color coded as indicated in the legend. As for the NLSF analysis (E, H), the larger dispersion of the phase lifetimes at lower concentrations is due to the increasing influence of the lower lifetime hotspots in the distribution. O: Scatter plot representation of data shown in E, H, illustrating the good agreement between ICCD measurements (horizontal axis) and SS2 measurements (vertical axis). P: Scatter plot representation of data shown in K, N, illustrating the good agreement between ICCD measurements (horizontal axis) and SS2 measurements (vertical axis). The dashed line indicates perfect agreement for O, P. Scale bar in A,B,D,J,K,M: 3 mm.

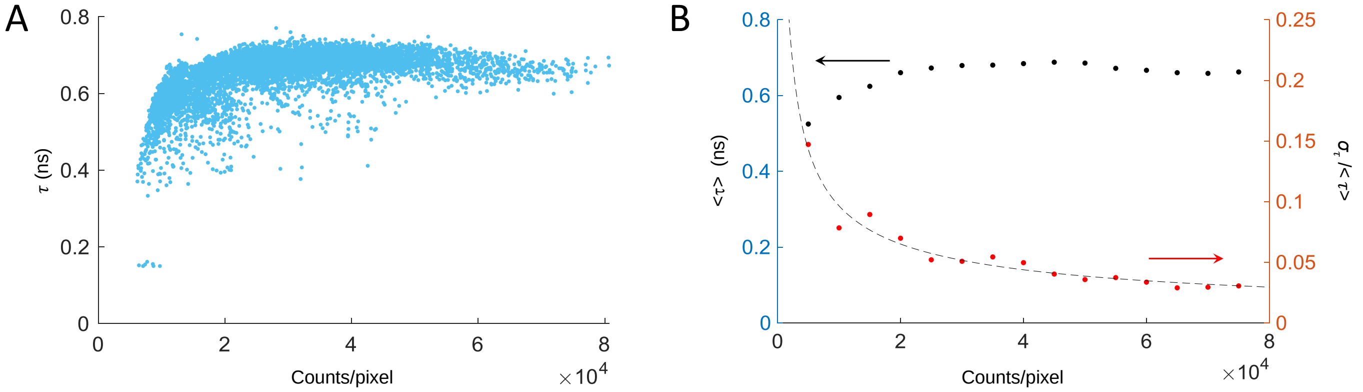

**Fig. S3**: Shot noise effect on lifetime precision. A: SS2 lifetime data corresponding to the sample discussed in Fig. S2D (single-exponential NLSF lifetime $\tau$) plotted as a function of pixel intensity $I$ (counts/pixel). B: Mean lifetime (black dots, left axis) and standard deviation (red dots, right axis) for each data slice $p/2 < 10^{-4} I < (p+1)/2$ in A, p = 0,…, 15. The standard deviation ($\sigma$) dependence on $I$ is well-fitted by a power law: $\sigma/\langle\tau\rangle = F I^{\beta}$ where $\beta = 0.568$, demonstrating a shot noise-limited behavior, with a F-factor $F = 18.05$.

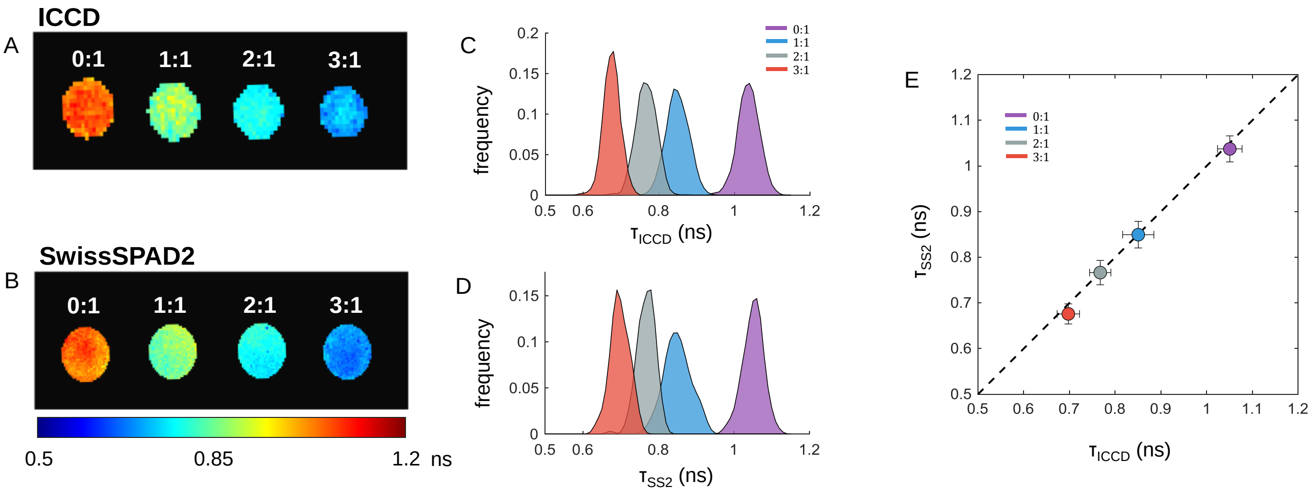

**Fig. S4**: NIR MFLI-FRET well-plate, single-exponential NLSF analysis. A, B: Single-exponential NLSF lifetime maps obtained for the ICCD and SS2. C, D: Fitted lifetime KDE distributions for each well, identified in the legend. E: Comparison between SS2-fitted lifetimes and ICCD-fitted lifetimes. The dashed line would indicate identity between the two measurements. The maximum relative deviation between SS2 and ICCD results for all samples is 1.1%.

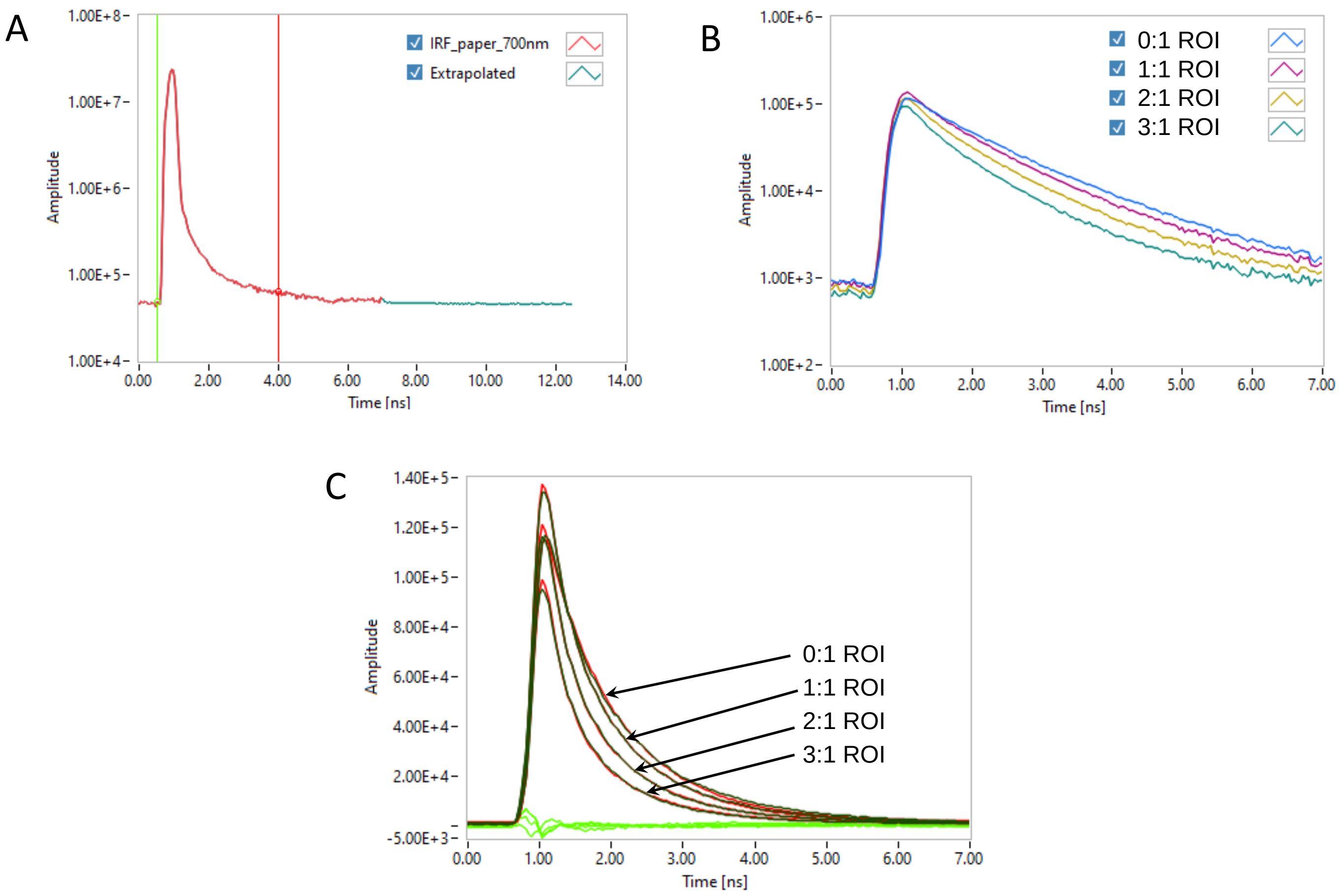

**Fig. S5**: Steps involved in obtaining global single-exponential references for bi-exponential NLSF analysis. A: Full-frame ICCD IRF decay (red curve) and its extrapolation (teal curve) to a full period duration (12.5 ns) used for decay convolution. B: Full-ROI fluorescence decays obtained from the dataset described in Fig. 2. Each ROI is identified in the legend. The donor-only decay (0:1) is clearly characterized by a longer lifetime than the others, which are decaying faster as the acceptor fraction increases. C: Result of bi-exponential fits for 1:1, 2:1 and 3:1 decays and single-exponential fit for the 0:1 decay (red curves) and residuals (green curves). The 0:1 decay was fitted without constraints, resulting in $\tau_1$ = 1 ns. This lifetime was used as constraint for one of the components of the bi-exponential fits performed on the other decays, resulting in the following second components: (1:1): $\tau_2$ = 0.273 ns, (2:1): $\tau_2$ = 0.264 ns, (3:1): $\tau_2$ = 0.257 ns. The mean value $\tau_2$ = 0.265 ns was therefore used as the second fixed component for the *single-pixel* bi-exponential NLSF analysis presented in Fig. 2.

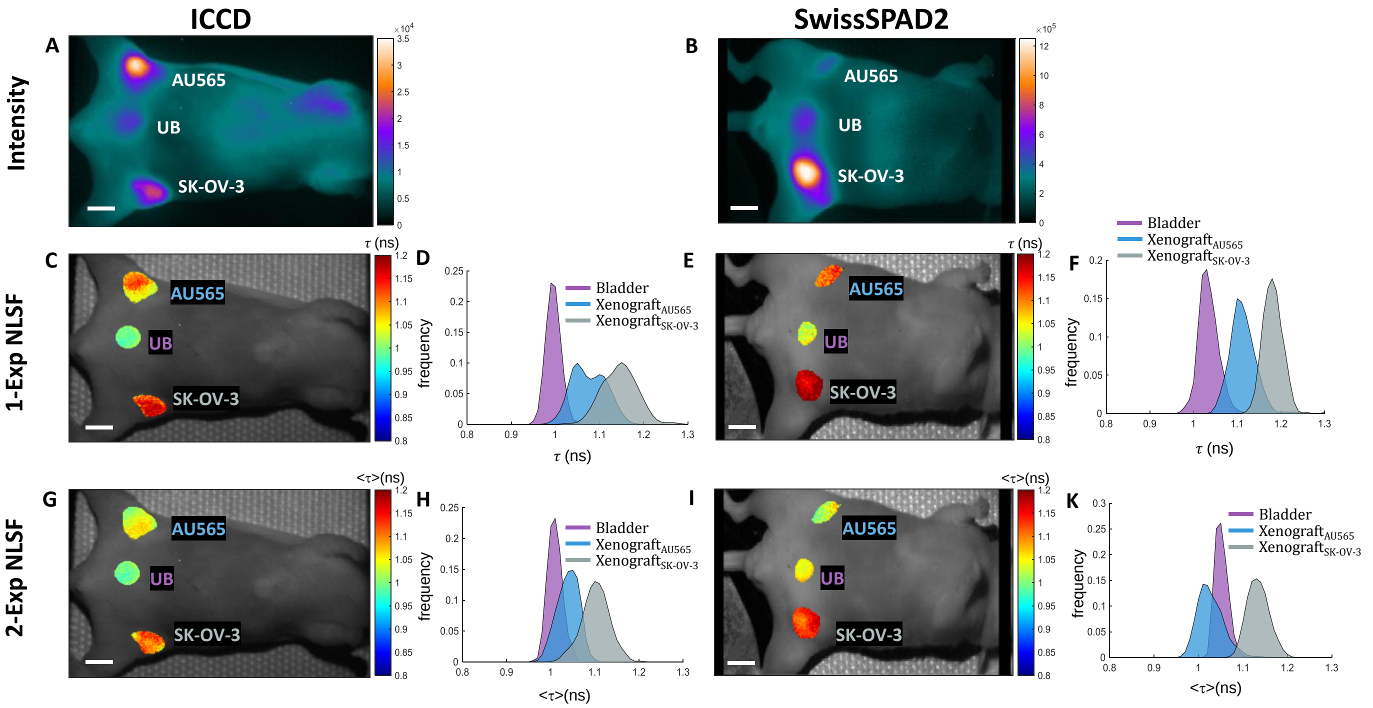

**Fig. S6**: MFLI-FRET TZM day 1: 1-exp vs average bi-exp NLSF lifetime. A, B: Whole body fluorescence intensity images for ICCD and SS2. C, E: Corresponding mono-exponential lifetime maps (calibrated via mouse reflectance, see *Methods*) for both tumors and the urinary bladder ROIs. D, F: 1-exponential lifetime KDE distributions obtained with both cameras. G, I: Corresponding 2-exponential lifetime maps (calibrated via mouse reflectance, see *Methods*) for both tumors and the urinary bladder ROIs. H, K: 2-exponential lifetime KDE distributions obtained with both cameras. Notice the similarity of the results of 1- and 2-exp NLSF for the UB and SK-OV-3, but lower average lifetimes obtained for AU565. Scale bar in A, B, C, E, G, I: 6 mm.

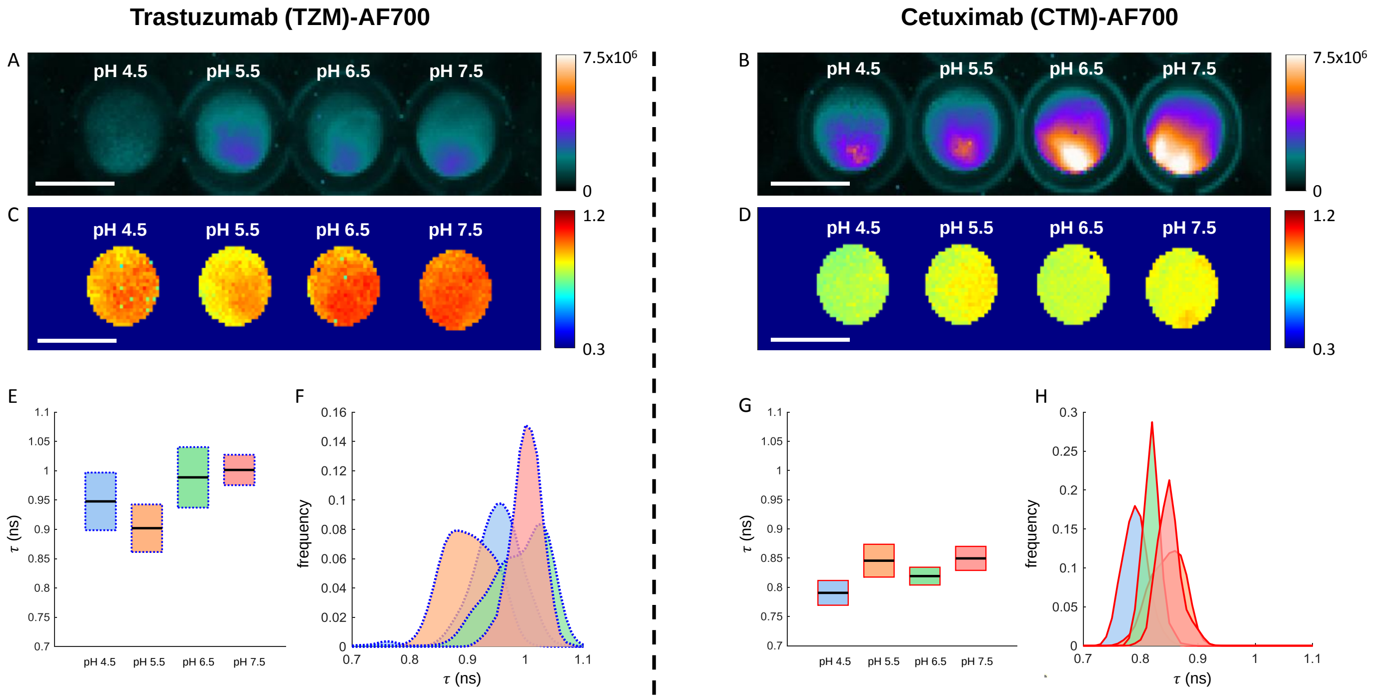

**Fig. S7**. Single-exponential NLSF analysis of SS2 acquisition of AF700 (NIR-FRET donor) conjugated to different molecules (Trastuzumab [TZM] and Cetuximab [CTM]) as a function of pH. A, B: Intensity images (in photon-counts) after background subtraction. C, D: Lifetime map retrieved by single-exponential NLSF analysis. E, G: Boxplot of lifetime as function of pH for both TZM (E) and CTM (G) samples. F: Lifetime distributions for all AF700-conjugated TZM wells. H: Lifetime distributions for all AF700-conjugated CTM wells.

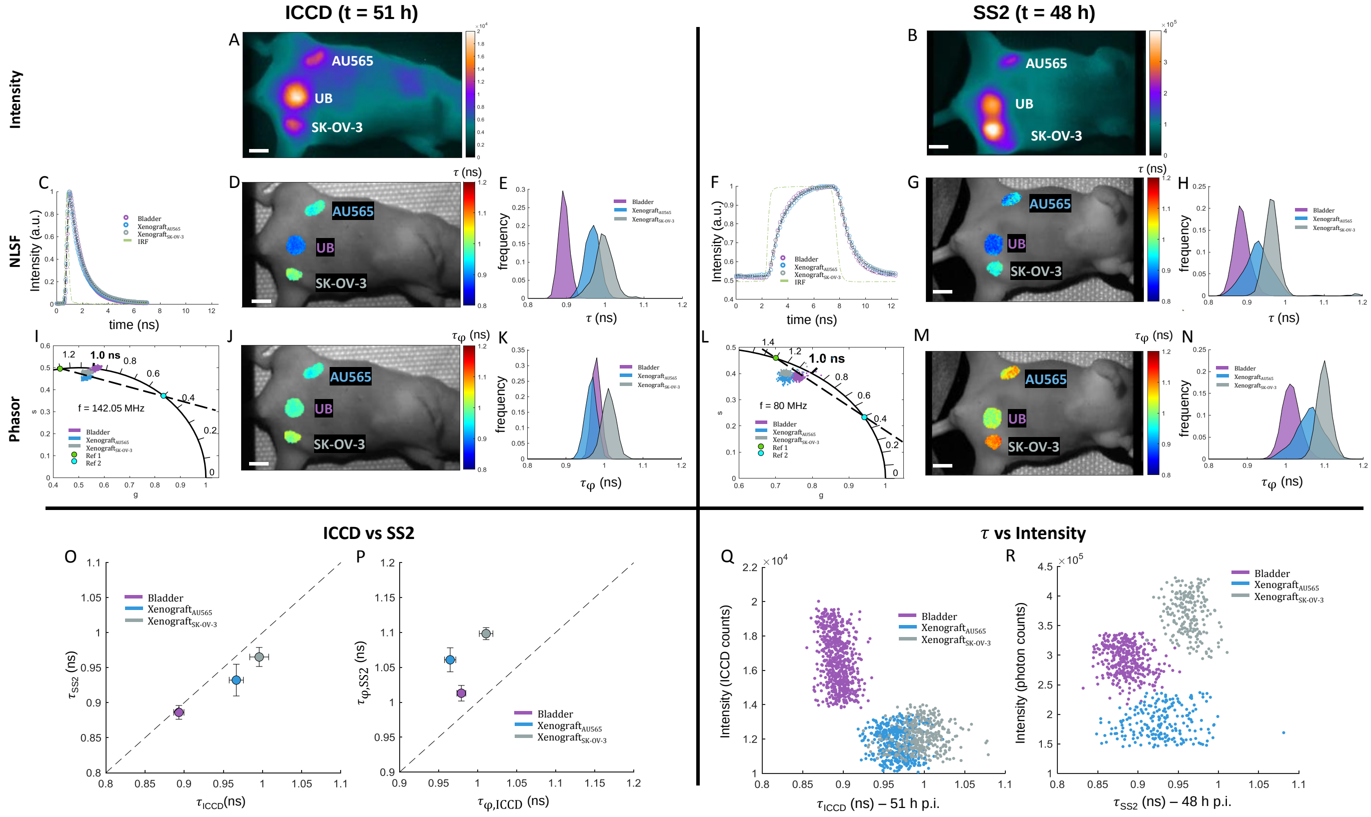

**Fig. S8:** MFLI-FRET TZM day 2: 1-Exp NLSF vs phase lifetime. One mouse was injected with 20 µg of AF700-TZM and 40 µg of AF750-TZM and imaged by MFLI at 48 h post-injection (p. i.) using the SS2 and 51 h post-injection using the ICCD. A, B: Fluorescence intensity images. A: ICCD; MCP voltage = 500 V, integration time/gate = 500 ms, illumination power = 2.13 mW/cm$^2$. B: SS2; integration time/gate = 2.65 s, illumination power = 3.2 mW/cm$^2$;. C, F: ICCD and SS2 normalized whole ROI decays for the different organs, plotted with the corresponding IRF. D, G: MFLI lifetime maps obtained by 1-exponential NLSF for both cameras. E, H: Corresponding lifetime KDE distributions for each ROI obtained for both cameras. I, L: phasor scatter plots color-coded by ROI, with overlaid reference lifetimes (green dot, reference 1; blue dot, reference 2) and dashed black line connecting them. J, M: pixel-wise phase lifetime maps and K, N: phase lifetime KDE distributions for the two xenografts. O: Scatter plot (mean ± standard deviation) showing the lifetime measured for each tumor with SS2 (48 h p.i.) versus that measured with the ICCD (same mouse, 51 h p.i.).  P: Scatter plot of phase lifetime results (mean ± standard deviation) for ICCD and SS2. Scale bar in A, B, D, G, K, N: 6 mm. Q, R: Scatter plots of intensity versus lifetimes retrieved via mono-exponential NLSF for the ICCD (Q) and SS2 (R). No correlation between fitted lifetime and intensity is observed.

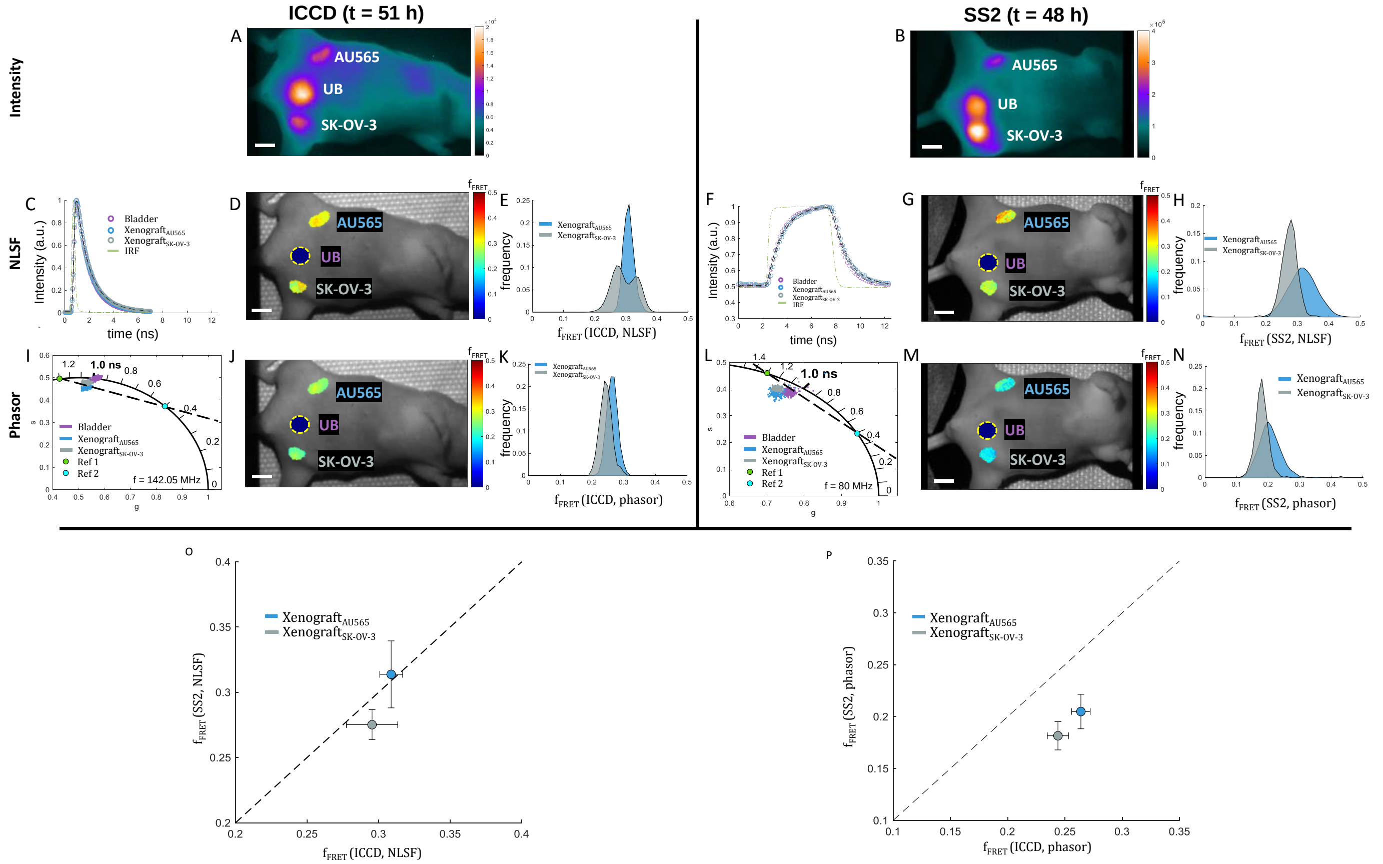

**Fig. S9:** MFLI-FRET TZM day 2: 2-Exp NLSF $f_{FRET}$ vs phasor ratio**.** One mouse was injected with 20 µg of AF700-TZM and 40 µg of AF750-TZM and imaged by MFLI at 48 h post-injection (p. i.) using the SS2 and 51 h post-injection using the ICCD. A, B: Fluorescence intensity images. A: ICCD; MCP voltage = 500 V, integration time/gate = 500 ms, illumination power = 2.13 mW/cm$^2$. B: SS2; integration time/gate = 2.65 s, illumination power = 3.2 mW/cm$^2$. C, F: ICCD and SS2 normalized whole ROI decays for the different organs, plotted with the corresponding IRF. D, G: MFLI FRET-fraction maps obtained by bi-exponential NLSF for both cameras. The urinary bladder (yellow dashed outline) was analyzed by 1-Exp NLSF and is therefore not included. E, H: Corresponding $f_{FRET}$ KDE distributions for each ROI obtained for both cameras I, L: phasor scatter plots color-coded by ROI, with overlaid reference lifetimes (green dot, reference 1; blue dot, reference 2) and dashed black line connecting them. J, M: pixel-wise phasor ratio maps and K, N: phasor-ratio KDE distributions for the two xenografts. O: Scatter plot (mean ± standard deviation) showing the FRET-fraction measured for each tumor with SS2 (mouse 2) versus that measured with the ICCD (mouse 1). P: Scatter plot of phasor ratio results (mean ± standard deviation) for SS2 vs ICCD. Scale bar in A, B, E, G, J, M: 6 mm.

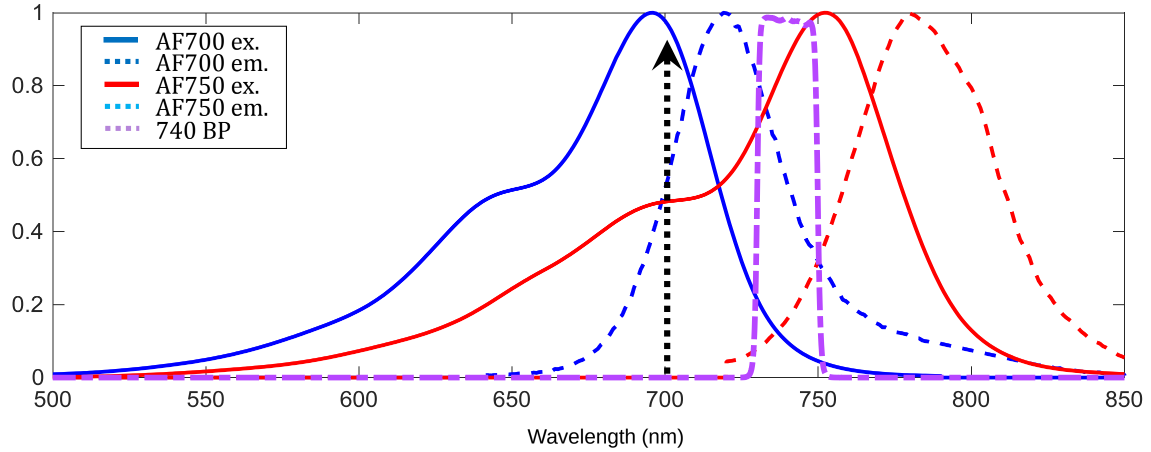

**Fig. S10**: Absorption and emission spectra of AF700/AF750 and bandpass emission filter. Spectral data obtained at https://www.chroma.com/spectra-viewer. Bandpass transmission data obtained at https://photonics.laser2000.co.uk/products/light-delivery-and-control/microscopy-filters/individual-filters/bandpass-filters/740-13-nm-brightline-single-band-bandpass-filter/ .

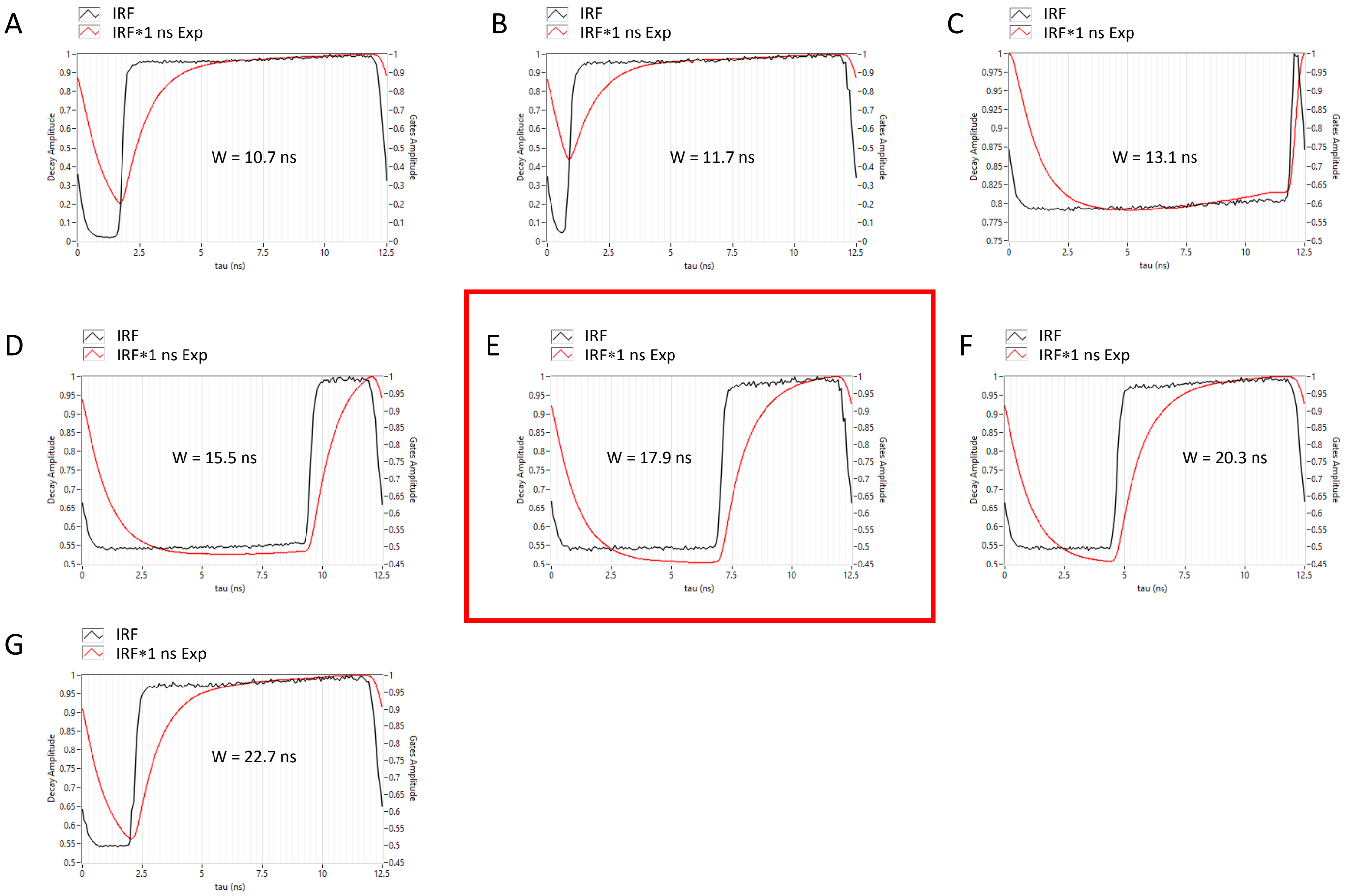

**Fig. S11**: Series of normalized paper IRFs (black curves, right vertical axis) obtained using the 7 available SS2 gate configurations, and the corresponding normalized convolution with a 1 ns single-exponential decay (red curves, left axis). For the shortest gate widths $W$, the decaying part of the red curve is truncated (it does not reach an asymptotic baseline), but the rising part is not (which is a symmetric version of the decaying part) and shows instead good convergence to an asymptote. For $W > T = 12.5$ ns, the situation is reversed until $W = 1.5\ T = 18.75$ ns, where rising and decaying parts both occupy half of the period. The gate width used in this work, $W$ = 17.9 ns is close enough from this situation, as is visible from the calculated decay in E.

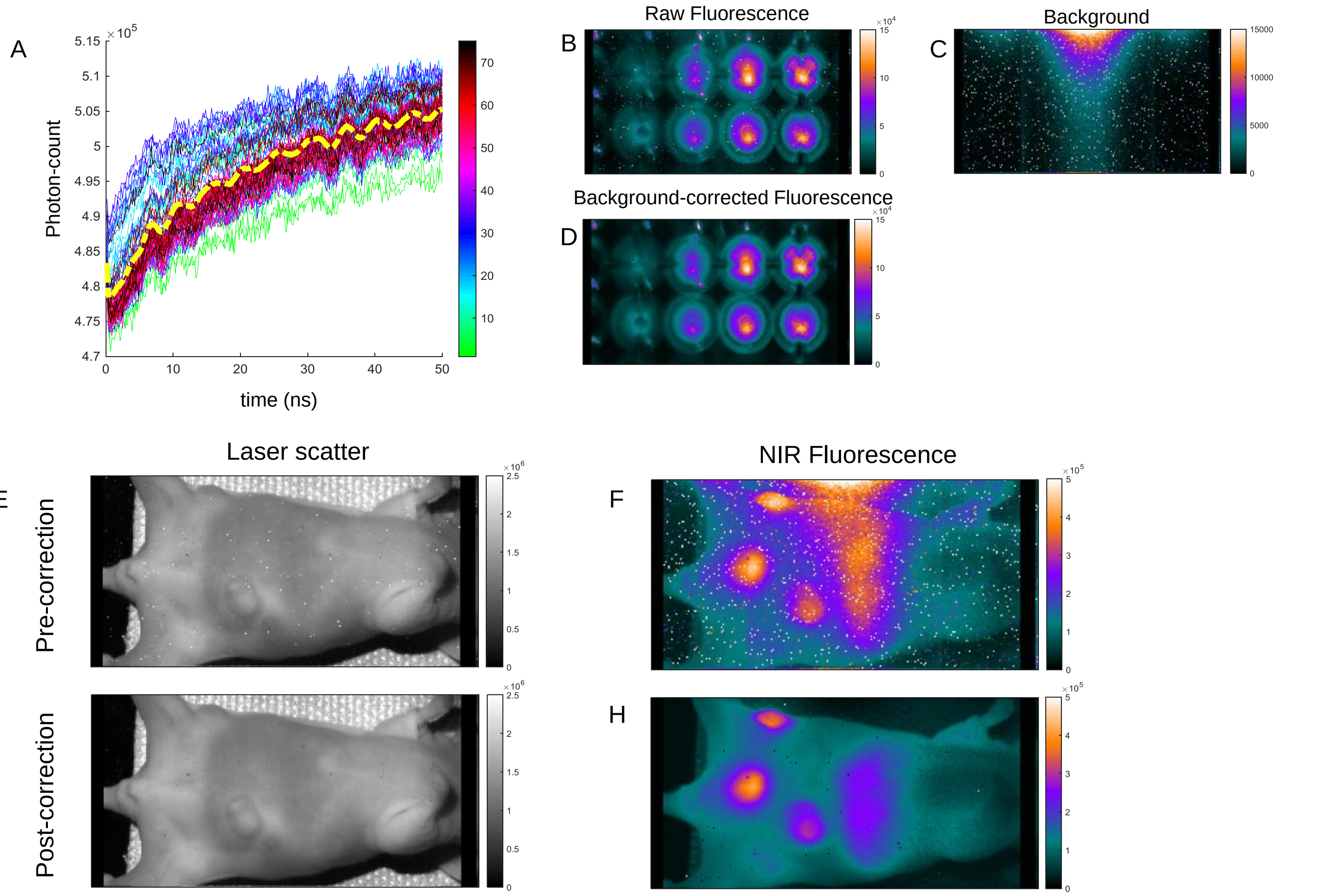

**Fig. S12**: SwissSPAD2 background correction workflow. A: SS2 background (summed intensity over all pixels) collected over a span of two hours. Each curve corresponds to one acquisition (each acquisition is separated from the next by 2 minutes) and is color coded by its location in the sequence as indicated by the color bar. Each curve is comprised of 280 gate values (0 – 50 ns) recording the background signal of the detector. A slight increase is observable from the first to the last gate acquisition, to which some small oscillations are superimposed. This non-flatness of the background "decay" can affect low-intensity sample decays if not corrected. The yellow dashed line indicates the averaged background profile, and is the result of small variations in detector response across time, likely due to temperature effects. B: uncorrected fluorescence, C: background signal and D: background-corrected fluorescence intensity images of the AF750 serial dilution well-plate experiment (Fig. S2). Color bars for B-D encode photon-count values. White spots in B, C correspond to noisy SPADs (or "screamers"), characterized by a high baseline count rate, which tend to dominate the fluorescence signal of actual sample, as seen in B. By subtracting C from B, the data shown in D is obtained, where most of these artifacts are eliminated. E-H: *in vivo* SS2 background correction. E: Uncorrected and G: background-corrected intensity image of the mouse studied in Fig. 4, acquired with 700 nm pulsed excitation without filter (scattered laser light used as IRF in the analysis). The main effect of background correction is to remove the "screamer" pixels (white in E). F: Uncorrected and H: background-corrected fluorescence intensity image (700 nm pulsed excitation and 740 bandpass emission filter) of the mouse studied in Fig. 4. In addition to removing most of the screamer pixel noise, background correction also takes care of the inhomogeneous detector background noise visible as a "smear" from a hot spot in the top center of the image in F. In this situation with low fluorescence signal level, the detector background dominates the actual signal in some areas of the detector (note the center top "halo" and the banded aspect of the background in F).

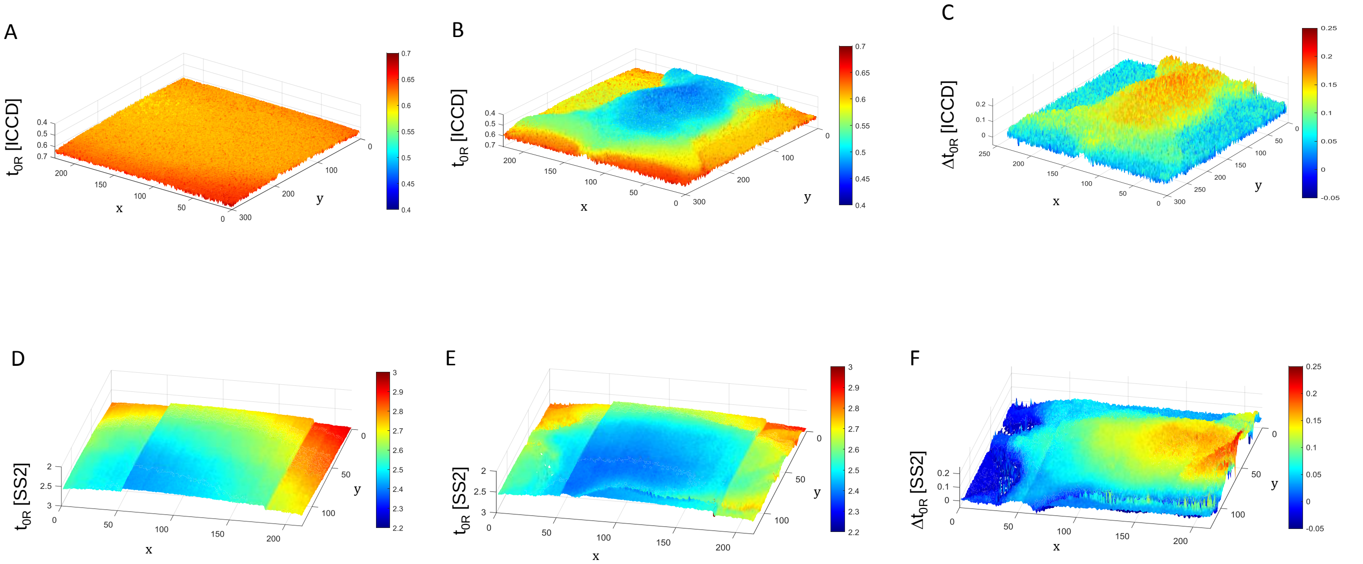

**Fig. S13**: A, B: Rising edge locations $t_{0R}$ retrieved from reflectance data with a white sheet of printer paper (A) and a mouse laying on a warming pad (B) with the ICCD. C: the difference, $\Delta t_{0R} = t_{0R}$ (paper) - $t_{0R}$ (mouse). The rising edge location is uniform for the paper sample but reveals a distinct topographical pattern in the mouse case, showing that the IRF delay is due to the path length difference between the surface of the mouse and the sample-supporting plane. D, E: equivalent data obtained with SS2. The non-uniformity of the detector's response is clearly visible in D, which shows three distinct rectangular regions, within which some additional variation is observable. The $t_{0R}$ map obtained with the mouse (E) is severely affected by this underlying pattern. F: the difference map E – D, which cancels out the intrinsic detector inhomogeneity reveals the same topographic pattern demonstrating again that the IRF delay is due to the path length difference between the surface of the mouse and the sample-supporting plane.

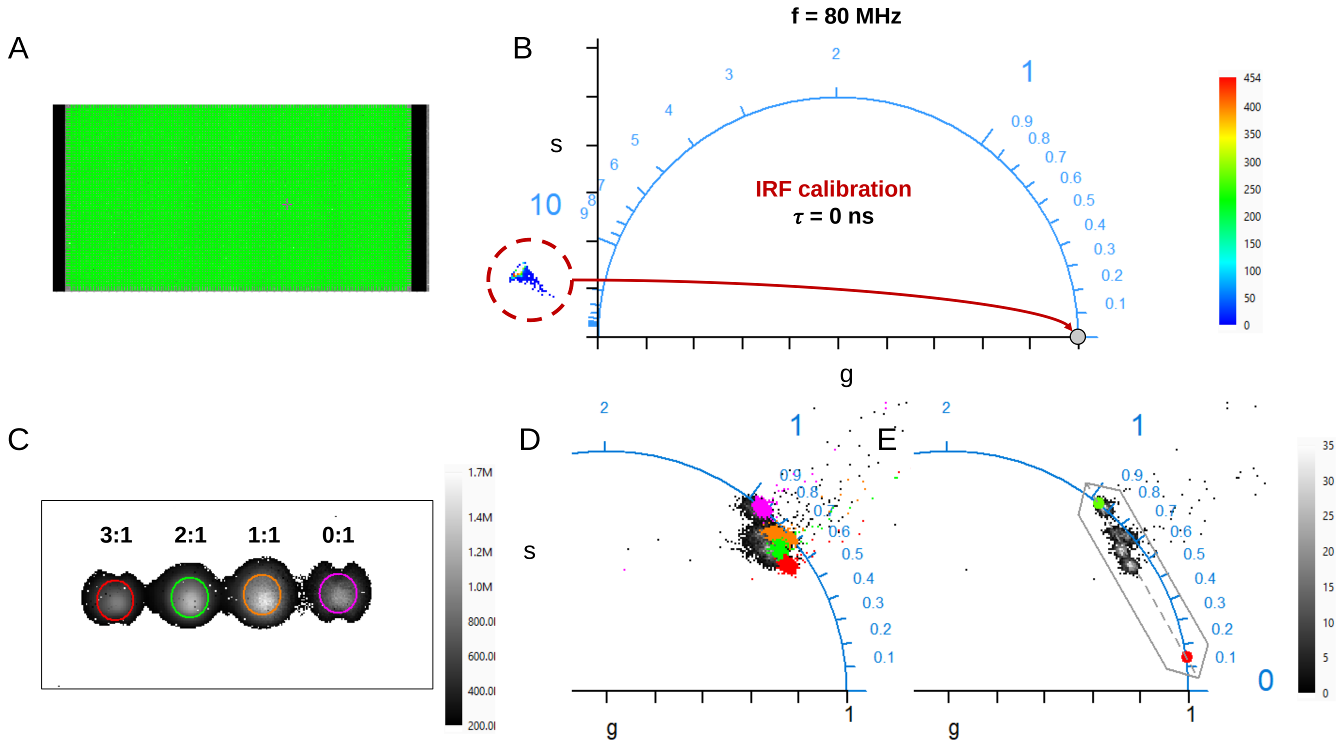

**Fig. S14**: Visual depiction of the phasor analysis workflow used for the NIR FRET well-plate SS2 data (Fig. 2). A: Map of pixel ROIs used for pixel-wise IRF dataset analysis. B: The histogram of uncorrected IRF phasors ($\tau$ = 0 ns) for all pixels shown in A are visible to the left of the universal circle (plotted in blue, with indication of the location of single-exponential phasors with ticks for 0, 0.1, …, 0.9, 1, 2, …, 10, etc. ns). The color bar on the right is used to encode the histogram bin values. After calibration using the IRF dataset, all these phasors end up at the (1, 0) point ($\tau$ = 0 ns) as indicated by the red arrow. C: Intensity image of the FRET sample well-plate, with 4 selected ROIs. Pixels with intensity < 200k counts are color-coded white. D: Highlighted calibrated phasors, the color indicating which ROI the phasors originate from. Phasors not highlighted correspond to pixels outside the selected ROIs E: Phasor plot limited to pixels within the selected ROIs (gray-scale histogram, color bar on the right common to D & E). The reference lifetimes used for phasor ratio analysis are indicated by a green and red dot, with the segment connecting them indicated by a dashed line. The IRF dataset was obtained using a white sheet of printer paper illuminated with the same source as the sample (700 nm pulsed laser), without the emission bandpass filter. Calibration lifetime was set to 0 ns, phasor frequency was set to f = 1/T = 80 MHz.

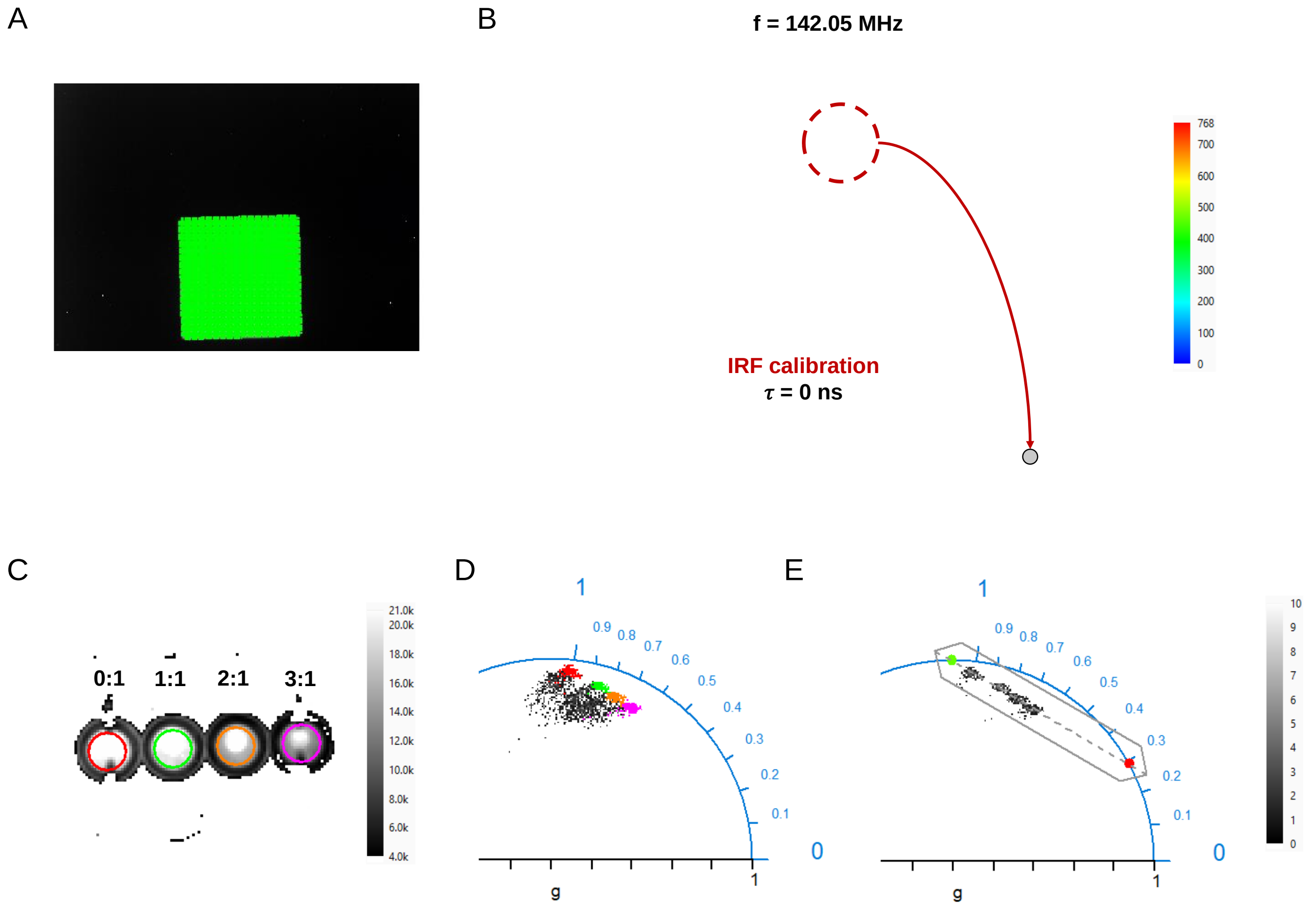

**Fig. S15**: Visual depiction of the phasor analysis workflow used for the *in vivo* CTM SS2 data (Fig. 4). A: Map of pixel ROIs used for pixel-wise IRF dataset analysis (mouse observed without emission filter). B: The histogram of uncorrected IRF phasors ($\tau$ = 0 ns) for all pixels in A are visible on the bottom left of the universal circle (plotted in blue, with indication of the location of single-exponential phasors with ticks for 0, 0.1, …, 0.9, 1, 2, …, 10, etc. ns). The color bar on the right is used to encode the histogram bin values. After calibration using the IRF dataset, the phasors of highlighted pixels end up at the (1, 0) point ($\tau$ = 0 ns) as indicated by the red arrow. C: Intensity image of the in vivo CTM dataset, with 4 selected ROIs. D: color-coded calibrated phasors, the color indicating which ROI the phasors originate from. Calibration lifetime: 0 ns, phasor frequency: 80 MHz.

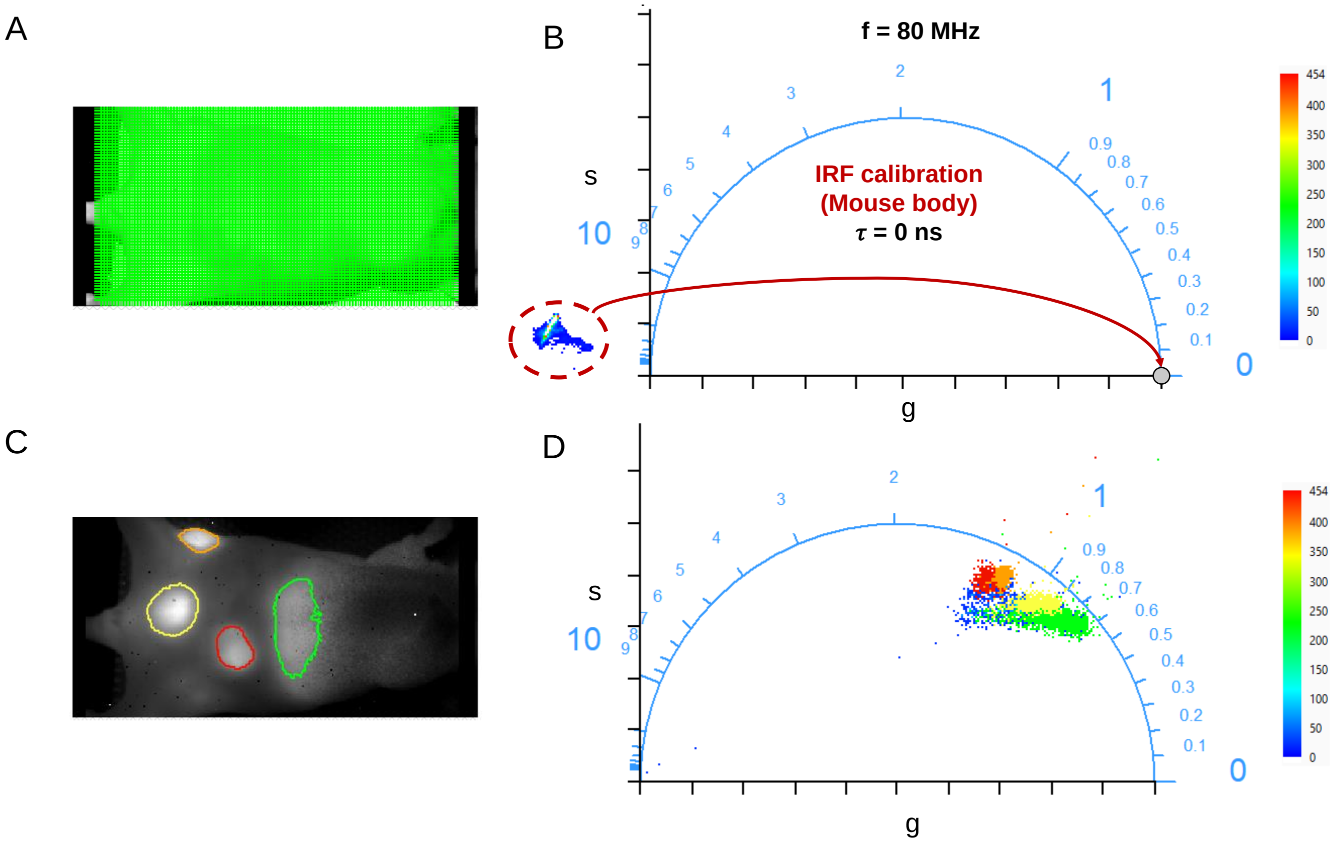

**Fig. S16:** Visual depiction of the phasor analysis workflow used for the NIR FRET well-plate ICCD data **(**Fig. 2). A: Map of pixel ROIs used for pixel-wise IRF dataset analysis. B: The histogram of uncorrected IRF phasors ($\tau$ = 0 ns) for all pixels in A are visible from the bottom left ((0, 0) point) to the top right of the universal circle (plotted in blue, with indication of the location of single-exponential phasors with ticks for 0, 0.1, …, 0.9, 1, 2, …, 10, etc. ns). The color bar on the right is used to encode the histogram bin values. The top-right cluster corresponds to the single-pixel ROIs highlighted in A, while the cluster around the origin corresponds to the remainder of the pixels, which correspond to uncorrelated detector noise. After calibration using the IRF dataset, the phasors of highlighted pixels end up at the (1, 0) point ($\tau$ = 0 ns) as indicated by the red arrow, while the other phasors remain in the vicinity of the origin. C: Intensity image of the FRET sample well-plate, with 4 selected ROIs. Pixels with intensity < 4k counts are color-coded white. D: Highlighted calibrated phasors, the color indicating which ROI the phasors originate from. Phasors not highlighted correspond to pixels outside the selected ROIs E: Phasor plot limited to pixels within the selected ROIs (gray-scale histogram, color bar on the right common to D & E). The reference lifetimes used for phasor ratio analysis are indicated by a green and red dot, with the segment connecting them indicated by a dashed line. The IRF dataset was obtained using a white sheet of printer paper illuminated with the same source as the sample (700 nm pulsed laser), without the emission bandpass filter. Calibration lifetime was set to 0 ns, phasor frequency was set to f = 1/D = 142.05 MHz.

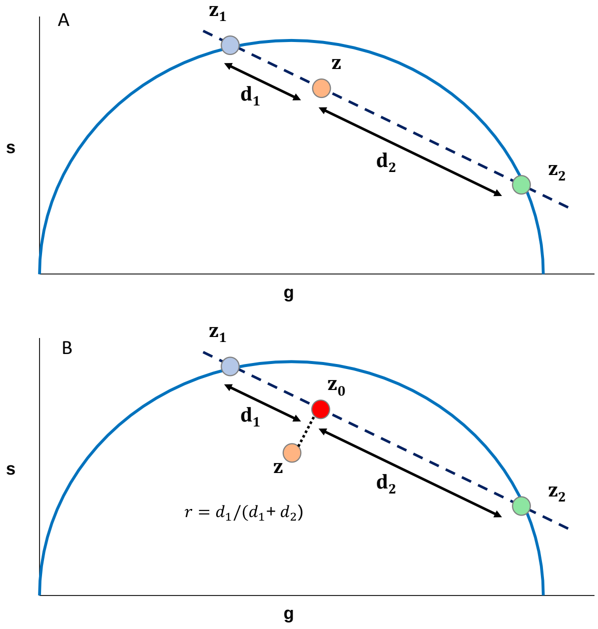

**Fig. S17**. Illustration of the principle of phasor ratio analysis in MFLI-FRET. A: In the ideal case, a mixture of two components with phasor $z_1$ and $z_2$ will be located on the segment (dashed line) connecting both components. The relative distances $d_1/d$ and $d_2/d$ (or phasor ratios, where $d = d_1 + d_2$) correspond to the fractions of each component $z_2$ or $z_1$ respectively. B: in the general case, a sample's phasor is not located exactly on the segment connecting both reference phasors. In those cases, the sample's phasor is first projected orthogonally to obtain $z_0$, which is used to compute the phasor ratios.

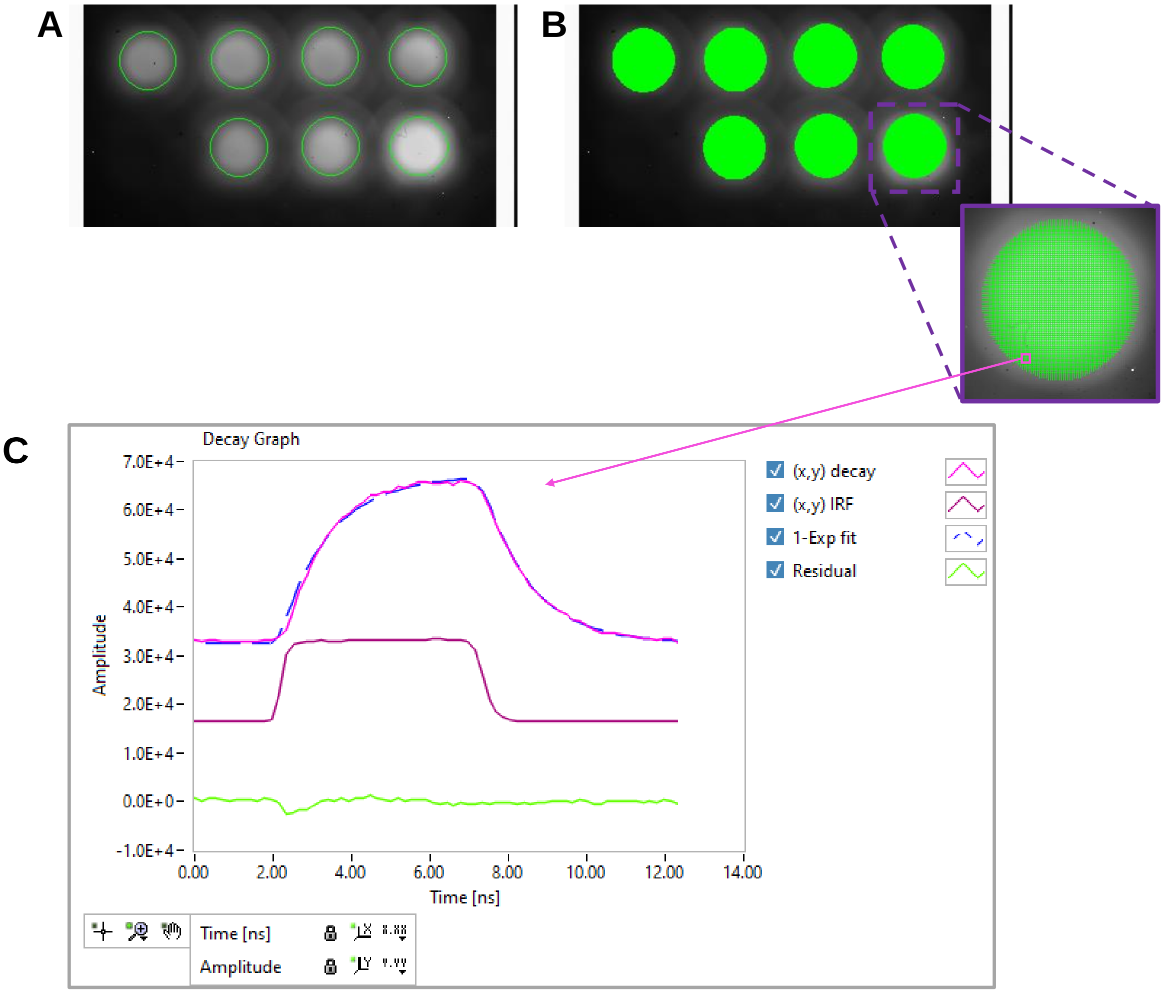

**Fig. S18**: Schematic depiction of the nonlinear least squares fit (NLSF) analysis workflow with Alligator. A: The intensity image (sum over all time-gates) is used to define regions of interest (ROIs) to which the analysis is limited. B: Each pixel in all ROIs is analyzed individually, with the corresponding pixel IRF decay used for convolution. C: Example of single-pixel analysis using a single-exponential model: the plot shows the original decay, together with the fitted curve (dashed blue curve) and the residuals (green curve). The corresponding local IRF used for convolution is also shown. In practice, all pixels are analyzed in batch mode and maps of each output parameters (including $\chi^2$ and $R^2$ coefficient) can then be examined graphically and exported for further analysis.

### Supplementary Notes

#### Supporting Note 1: NIR-MFLI lifetime measurement as a function of concentration (Alexa Fluor 750)

One of the advantages of lifetime contrast is its independence on dye intensity or concentration. However, this independence is limited by dye photophysics (e.g. dye saturation at high excitation intensities, or self-absorption at high concentration) and detector physics (including pile-up due to detector deadtime at high intensities [4]). To investigate SS2's ability to perform accurate and quantitative NIR-MFLI independently of intensity or concentration, we imaged solutions of Alexa Fluor 750 (AF750) in water or phosphate buffer saline (PBS) at different concentrations. AF750 is a NIR fluorescent dye emitting at wavelengths $\lambda > 750$ nm and characterized by a lifetime $\tau = 0.65$ ns at pH 7 [5]. This a priori makes AF750 a challenging benchmark for SS2, whose gate rise time (640 ps) is of the same order of magnitude as AF750's lifetime [1].

##### NLSF analysis

Fluorescence intensity images as well as representative single-pixel decays and corresponding NLSF results and residuals (see *Methods*) are shown in Fig. 1A, B and Fig. 1C, F, respectively. These curves illustrate the different aspects of ICCD and SS2 data, and excellent fits are generated in both cases. The lifetime maps obtained with both cameras are shown in Fig. S2D and Fig. 2G, respectively. Despite the detector differences and the fact that data was acquired on two distinct setups, the lifetime maps obtained with both detectors are in good agreement with one another (Fig. S2D, G), with a maximum difference of 4.5% between the lifetimes measured by the two detectors (Fig. S2O). Interestingly, the presence of caustics (laser excitation light concentrated in specific locations within the wells), which are clearly visible in the intensity images, does not affect the lifetime map uniformity at *high* concentration but results in "hotspots" with shorter lifetimes at lower concentrations. These "hotspots" exhibiting lower lifetimes are probably due to contamination by autofluorescence of the well material, which, should become more prominent at lower dye concentrations. These results demonstrate the ability of SS2 to extract NIR lifetimes below 1 ns independently of signal intensity (in the absence of background contamination). Moreover, lifetime variance as a function of signal intensity is inversely proportional to the signal (Supplementary Note 1 & Supplementary Fig. S3), confirming that these measurements are shot noise limited[6], [7].

To verify the shot noise-limited behavior of the NLSF analysis of SS2 data, the fitted-lifetime data corresponding to the sample discussed in Fig. S2D (single-exponential NLSF lifetime $\tau$) was plotted as a function of pixel intensity $I$ (Fig. S3A) Slicing the data into intervals defined by $p/2 < 10^{-4} I < (p+1)/2$, we computed the mean and standard deviation of the lifetime in each slice, obtaining two curves shown in Fig. S3B. The standard deviation's dependence on $I$ is well-fitted by a power law: $\sigma/<\tau> = F I^{-\beta}$ where $\beta = 0.501$, demonstrating a shot noise-limited behavior, with an F-factor $F = 18.05$. This result is in good agreement with the values obtained for visible range dyes in ref. [3] ($F \approx 4$). The mean lifetime itself is characterized by a negative bias at low intensity values, which is due to the contamination by well autofluorescence, characterized by a shorter lifetime, as discussed in the main text.

##### Phasor analysis

Partial phasor plots for each well (Fig. S2I, L), as well as corresponding pixel-wise phase lifetime maps (Fig. S2J, M), are shown for the ICCD and SS2 data. Since ICCD data covered a duration $D = 10.04$ ns smaller than the laser period, $T = 12.5$ ns, a phasor frequency $f = 1/D = 99.6$ MHz was used, while the SS2 data covering the whole laser period, $f = 1/T = 80$ MHz was used. As expected, the phase lifetime maps obtained with both cameras are in good agreement with one another, a result better illustrated by the boxplot representations of Fig. S2K, N. The maximum relative difference of 5.9% between phase lifetimes measured with the ICCD or SS2 (Fig. S2P) is comparable to the NLSF results. More importantly, the NLSF-computed lifetimes and phase lifetimes are in excellent agreement with one another (Supplementary Table 1), even considering the probable imperfection of the paper IRF used for decay convolution (NSLF) and phasor calibration (phasor analysis). This analysis establishes the ability of both detector technologies, using two distinct FLI analysis approaches, to quantitatively resolve sub-nanosecond lifetimes of a NIR dye at low concentrations, setting the main foundation for the remainder of this study.

#### Supplementary Note 2: Further discussion of Day 2 *in vivo* TZM data

Fig. S8A, B shows the resulting fluorescence intensity maps obtained three hours apart. Three bright regions are clearly visible: the two tumor xenografts (top: SK-OV-3, bottom: AU565) and the urinary bladder (center: UB). As an excretion organ, the signal originating in the urinary bladder probably indicates degradation products, as discussed in prior work[6]. Therefore, bladder oftentimes displays variable AF700-TZM lifetimes from animal to animal, or from one time point to the next within a single animal[6], [7]. As is apparent in Fig. S8A, B, a comparable variation of the relative intensities of the three different regions (tumors and urinary bladder) is observed across time: the SK-OV-3 tumor shows the largest fluorescence level at 48 h (Fig. S8B, SS2 imaging), similar to that in the bladder at that time point, but drops to the level of the AU565 tumor 3 h later (Fig. S8A, ICCD imaging). While fluorescence photobleaching may play a role in these intensity variations, it cannot fully account for them, in particular their heterogeneity. Differences in relative probe accumulation over time could also contribute to these observations, but so could many other contributing factors, exemplifying the difficulties involved with using intensity alone to analyze molecular interactions by FRET *in vivo*.

Despite the intensity variations, single-exponential NLSF analysis of the time-resolved data (Fig. S8C, F**)** provides a more consistent picture within each ROI. Lifetimes measured 3 h apart with one or the other camera are similar between xenografts and across time, despite some systematically longer lifetimes reported in the later ICCD measurement (compare Fig. S8D, E and S6C, E). In particular, SK-OV-3 appears to consistently show the longest lifetime, despite a large relative intensity change, while the bladder is consistently characterized by the shortest lifetimes. As expected, no correlation between lifetime and relative intensity appears to exist (SI Fig. S8Q, R), supporting the use of MFLI as a complement to intensity information for any *in vivo* molecular imaging experiment.

These single-exponential NSLF results are confirmed by phasor analysis of the same datasets (Fig. S8I-N, P), revealing the same similar AF700 lifetimes in all ROIs, with small relative differences in general agreement with those reported by NLSF analysis. This agreement is all the more remarkable that in both cases (as discussed in *Methods* and *Supplementary Note* 5), the dataset used as IRF for decay convolution in NLSF analysis, and for phasor calibration in phasor analysis, is at best an approximation to the true IRF, which is inaccessible in this simple MFLI approach.

#### Supplementary Note 3: Decay convolution with SS2's IRF at repetition rates $f$ = 80 MHz

SS2 available gate durations are $W$ = 10.7, 11.7, 13.1, 15.5, 17.9, 20.3 or 22.7 ns. To visualize the effect of these gate durations on a single-exponential decay, we computed the convolution of the measured IRF (obtained by imaging the laser light scattered by a sheet of white filter paper) with a theoretical 1 ns single-exponential decay using Phasor Explorer, the software introduced in ref. [8]. The corresponding normalized curves are shown on Fig. S11, with the IRF shown in black (right vertical axis) and the convolved decay in red (left vertical axis).

The IRF width which provides a similar range for both the decaying part of the convolved curve and its rising part, is indicated by the red frame and corresponds to $W$ = 17.9 ns (sometimes referred to as config 5 in the raw data file names found on the Figshare repository). An acceptable alternative would have been $W$ = 20.3 ns.

Note that the shortest gate ($W$ = 10.7 ns) results in a fairly complete rising part (in the sense that the curve reaches an asymptote after the initial rise), but in a truncated decaying part. This effect is increasing until $W$ = 13.1 ns, where the *effective* width of the IRF is 13.1 – 12.5 = 0.6 ns. However, because of the lack of flatness of the IRF baseline (visible in Fig. S11C), the resulting convolved decay does not look particularly similar to a standard TCSPC decay and was not retained for this work.

It is worth noting that for $W > T$ = 12.5 ns, decay and IRF are offset vertically due to a full period integration (note the different scale ranges).

#### Supplementary Note 4: AlliGator data analysis workflow

##### SS2 data preprocessing

Because each pixel is an independent detector characterized by individual noise properties, it is important to individually correct raw data. The first effect corrected for is pile-up [3], resulting from the finite storage capacity of the 1-bit counter associated with each SPAD. Next, detector background needs to be subtracted as detailed in Fig. S12, based on a detector background dataset (itself pile-up corrected, and generally averaged over many acquisitions to reduce statistical noise). Here, by contrast to what was done in previous works (e.g. [3] & [9]), where ROI-based correction was performed, we used pixel-level correction to allow optimal subsequent pixel-wise analysis.

##### NLSF

NLSF analysis in AlliGator can be performed at the ROI level or at the pixel level, either interactively or in batch mode. As illustrated in Fig. S18A, B, the intensity image (sum over all time-gates) is first used to define regions of interest (ROIs) to which the analysis is limited. Subsequently, each pixel in all selected ROIs is analyzed individually, with the corresponding pixel IRF decay (obtained from the appropriate dataset) used for convolution. Fig. S18C shows an example of interactive single-pixel analysis using a single-exponential model: the plot shows the original decay, together with the fitted curve (dashed blue curve) and the residuals (green curve). The corresponding local IRF used for convolution is also shown.

In practice, all pixels are analyzed in batch mode and maps of each output parameters (including $\chi^2$ and $R^2$ coefficient) can then be examined graphically and exported for further analysis.

In the case of ICCD data, which is generally not covering the whole laser period, the IRF needs to be extrapolated to extend it up to the end of the period. This is done automatically by assuming that the tail of the IRF (missing decaying part) is exponential and asymptotically matches the recorded IRF at the beginning of the period (the IRF, like the sample decay, is periodic by definition).

#### Supplementary Note 5: Mouse IRF equivalent topographic maps

In order to visually illustrate the difference between the scattered laser path in the *in vitro* (paper) and *in vivo* (mouse) cases, we calculated the IRF's rising edge location at each pixel of interest for both paper and mouse IRF data files, by fitting a piece-wise logistic square gate model $f(t)$ to the recorded IRF:

$$\begin{cases} t \in [t_{R1}, t_{R2}]: & f_R(t) = A_1 - \dfrac{H_1}{1 + \exp\left(\left(t - t_{0,R}\right)/\sigma_R\right)} \\ t \in [t_{R2}, t_{F1}]: & f_U(t) = A_H \\ t \in [t_{F1}, t_{F2}]: & f_F(t) = A_2 - H_2 + \dfrac{H_2}{1 + \exp\left(\left(t - t_{0,F}\right)/\sigma_F\right)} \\ t \in [t_{start}, t_{R1}] \cup [t_{F2}, t_{end}]: & f_L(t) = A_L \end{cases} \tag{S1}$$

where the first interval brackets correspond to the rising edge, the second to the gate "on" state, the third brackets to the falling edge, and the last two correspond to the gate "off" state. $t_{0,R}$ (resp. $t_{0,F}$) represents the location of the rising edge midpoint and $\sigma_R$ (resp. $\sigma_F$) a measure of its width. The other parameters are obtained by imposing continuity between the different intervals of the model, the boundaries being graphically defined by the user. This model (or a variant with tilted "on" state) is a good approximation of the SS2 gate profile, and an acceptable one for the ICCD IRF (using $t_{R2} = t_{F1}$). Since we are only using it to illustrate differences between paper and mouse IRF, a detailed discussion of this model will be provided elsewhere.

As shown in Fig. S13A, in the case of the ICCD, the paper IRF's rise time locations are uniform over the whole field of view except for a slight curvature which might reflect the MCP response non-uniformity. By contrast, the mouse IRF rising edge location map (B) is strikingly similar to a topographic map of the mouse surface with delays between the "earliest" IRF and the "latest" one on the order of 150 ps, corresponding to ~45 mm = 2×22.5 mm of light propagation difference, consistent with a mouse height of ~ 2 cm.

In the case of SS2, the paper IRF delay map (D) shows a characteristic "band" pattern due to electronic signal routing differences[1], which are also observed in the SS2 mouse IRF dataset (E). Subtracting the former from the latter (F) removes the detector contribution to the rising edge location, resulting in a map also reminiscent of the mouse surface topography, with a similar delay range as that observed in the ICCD case.
